# Structure of herpes simplex virus pUL7:pUL51, a conserved complex required for efficient herpesvirus assembly

**DOI:** 10.1101/810663

**Authors:** Benjamin G. Butt, Danielle J. Owen, Cy M. Jeffries, Lyudmila Ivanova, Jack W. Houghton, Md. Firoz Ahmed, Robin Antrobus, Dmitri I. Svergun, John J. Welch, Colin M. Crump, Stephen C. Graham

**Author notes:** Corresponding author: Stephen C Graham. **Author Contributions** Conceptualization: CMC, SCG; Data curation: BGB, SCG; Formal Analysis: JJW; Funding Acquisition: DIS, CMC, SCG; Investigation: BGB, DJO, CMJ, LI, MFA, JWH, SCG; Methodology: CMJ, RA, DIS, JJW; Project Administration: SCG; Resources: CMC, SCG; Software: BGB, SCG, JJW; Supervision: SCG; Validation: BGB, MFA; Visualization: BGB, SCG; Writing – original draft: BGB, SCG; Writing – review & editing: All authors.

## Abstract

Herpesviruses are an ancient family of highly-prevalent human and animal pathogens that acquire their membrane envelopes in the cytoplasm of infected cells. While multiple conserved viral proteins are known to be required for efficient herpesvirus production, many of these proteins lack identifiable structural homologues and the molecular details of herpesvirus assembly remain unclear. We have characterized the complex of assembly proteins pUL7 and pUL51 from herpes simplex virus (HSV)-1, an α-herpesvirus, using multi-angle light scattering and small-angle X-ray scattering with chemical crosslinking. HSV-1 pUL7 and pUL51 form a stable 1:2 complex that is capable of higher-order oligomerization in solution. We solved the crystal structure of this complex, revealing a core heterodimer comprising pUL7 bound to residues 41–125 of pUL51. While pUL7 adopts a previously-unseen compact fold, the extended helix-turn-helix conformation of pUL51 resembles the cellular endosomal complex required for transport (ESCRT)-III component CHMP4B, suggesting a direct role for pUL51 in promoting membrane scission during virus assembly. We demonstrate that the interaction between pUL7 and pUL51 homologues is conserved across human α-, β- and γ-herpesviruses, as is their association with *trans*-Golgi membranes in cultured cells. However, pUL7 and pUL51 homologues do not form complexes with their respective partners from different virus families, suggesting that the molecular details of the interaction interface have diverged. Our results demonstrate that the pUL7:pUL51 complex is conserved across the herpesviruses and provide a structural framework for understanding its role in herpesvirus assembly.

**Significance Statement:** Herpesviruses are extremely common human pathogens that cause diseases ranging from cold sores to cancer. Herpesvirus acquire their membrane envelope in the cytoplasm via a conserved pathway, the molecular details of which remain unclear. We have solved the structure of a complex between herpes simplex virus (HSV)-1 proteins pUL7 and pUL51, two proteins that are required for efficient HSV-1 assembly. We show that formation of this complex is conserved across distantly-related human herpesviruses, as is the association of these homologues with cellular membranes that are used for virion assembly. While pUL7 adopts a previously-unseen fold, pUL51 resembles key cellular membrane-remodeling complex components, suggesting that the pUL7:pUL51 complex may play a direct role in deforming membranes to promote virion assembly.

## Introduction

Herpesviruses are highly prevalent human and animal pathogens that cause life-long infections and result in diseases ranging from cold sores and genital lesions (herpes simplex virus, HSV) to viral encephalitis (HSV-1), congenital birth defects (human cytomegalovirus, HCMV) and cancer (e.g. Kaposi’s sarcoma associated herpesvirus, KSHV) (1, 2). Herpesviruses share conserved virion morphology, their DNA genome-containing capsids being linked to glycoprotein-studded limiting membranes via a proteinaceous layer called *tegument*, and a conserved assembly pathway whereby final envelopment of the DNA-containing capsids occurs in the cytoplasm (reviewed in (3, 4)). While herpesviruses are known to extensively remodel the intracellular architecture of infected cells (5), the molecular mechanisms by which they direct intracellular membranes to envelop nascent virions remain unclear.

HSV-1 tegument proteins pUL7 and pUL51 form a complex that promotes virus assembly by stimulating the cytoplasmic wrapping of nascent virions (6, 7). pUL7 and pUL51 co-localize with Golgi markers both during infection and when co-transfected into cells (6–8), palmitoylation of residue Cys9 being required for pUL51 membrane association (8). Deletion of pUL7, pUL51, or both proteins from HSV-1 causes a 5- to 100-fold decrease in virus replication (6, 9, 10) and cells infected with HSV-1 lacking pUL7 and pUL51 accumulate unenveloped capsids in the cytoplasm (6). Similar results have been observed in other α-herpesviruses. pORF53 and pORF7, the pUL7 and pUL51 homologues from varicella-zoster virus (VZV), co-localize with *trans*-Golgi markers in infected cells (11, 12) and deletion of pORF7 causes a defect in cytoplasmic envelopment (13). Similarly, deletion of pUL7 or pUL51 from pseudorabies virus (PrV) causes defects in virus replication and the accumulation of cytoplasmic unenveloped virions (14, 15), and PrV pUL51 co-localizes with Golgi membranes during infection (14).

Homologues of pUL7 and pUL51 can be identified in β- and γ-herpesviruses, although pUL51 homologues lack significant sequence similarity with α-herpesvirus pUL51 and their homology is inferred from their conserved positions in virus genomes (16, 17). The putative pUL51 homologue pUL71 from HCMV, a β-herpesvirus, associates with the Golgi compartment when expressed in isolation and with Golgi-derived virus assembly compartments during infection (18). Deletion of pUL71 causes defects in HCMV replication, characterized by aberrant virus assembly compartments (19) and defects in secondary envelopment (20). Similarly, the HCMV pUL7 homologue pUL103 co-localizes with Golgi markers when expressed alone or during infection, and deletion of pUL103 causes a loss of assembly compartments, reductions in virus assembly and defects in secondary envelopment (21). Relatively little is known about the pUL7 and pUL51 homologues from γ-herpesviruses. Both the pUL7 and pUL51 homologues from murine γ-herpesvirus 68 are essential for virus replication (22). The putative pUL51 homologue BSRF1 from Epstein-Barr virus associates with Golgi membranes and siRNA knock-down of BSRF1 in B95-8 cells prevents virion production (23). The KSHV homologue of pUL7, pORF42, is similarly required for efficient virion production (24). While a direct interaction has not been shown for the pUL7 and pUL51 homologues from β- or γ-herpesviruses, the EBV homologues BBRF2 and BSRF1 have been shown to co-precipitate from transfected cells (23).

Definitive molecular characterization of pUL7 and pUL51 function in HSV-1 or other herpesviruses has been hampered by their lack of homology to any proteins of known structure or function. pUL7 binds directly to pUL51 and the abundance of either protein in infected cells is dramatically reduced when the other is absent, suggesting that they form an intimate complex that is required for their correct folding (6). We characterized the pUL7:pUL51 complex by solution scattering and solved the atomic-resolution structure of the proteolysis-resistant core of this complex using X-ray crystallography. pUL7 comprises a single globular domain that binds one molecule of pUL51 via a hydrophobic surface, a second molecule of pUL51 being recruited to the solution complex via the N-terminal region of pUL51. While the fold of pUL7 is not similar to any known structure, the α-helical pUL51 protein shares unanticipated structural similarity to components of the endosomal sorting complex required for transport (ESCRT)-III membrane-remodeling machinery. Furthermore, we show that formation of the pUL7:pUL51 complex and its association with the *trans*-Golgi network is conserved across α-, β- and γ-herpesviruses, consistent with a conserved function for this complex in herpesvirus assembly.

## Results

### HSV-1 pUL7 and pUL51 form a 1:2 heterotrimer in solution

Full-length HSV-1 pUL7 and pUL51 were co-expressed in *Escherichia coli*, the palmitoylation site of pUL51 (Cys9) having been mutated to serine to avoid aberrant disulfide bond formation (Fig. S1). Co-expression and co-purification was used because pUL51 formed large soluble aggregates when purified alone (Fig. S1) and pUL7 was extremely prone to aggregation upon removal of the GST purification tag when purified in the absence of pUL51. Multi-angle light scattering (MALS) analysis showed the complex to elute from size-exclusion chromatography (SEC) as two peaks with molecular masses of 79.0 ± 1.8 kDa and 165.5 ± 1.1 kDa (Fig. 1*A*), consistent with pUL7 and pUL51 forming a 1:2 heterotrimer in solution (calculated mass from amino acid sequence 84.5 kDa) that dimerizes at higher concentrations to form a 2:4 heterohexamer (calculated mass 169 kDa). However, pUL51 of the co-purified complex was prone to degradation, frustrating crystallization attempts (Fig. 1*A*). Prior sequence analysis (8, 25) and our bioinformatics (Fig. S2) suggested that the C-terminal region of pUL51 lacks regular secondary structural elements and is disordered. Consistent with this prediction, SEC with inline small-angle X-ray scattering (SAXS) showed the pUL7:pUL51 complex to be extended. The 1:2 and 2:4 complexes have radii of gyration (R_g_) of 4.3 and 4.8 nm, with maximum particle dimensions (*D*_*max*_) of ~18 nm and 20 nm, respectively (Fig. 1*B*, 1*J*, 1*K* and Table S1). *Ab initio* shape analysis was performed by fitting the 2:4 scattering curve to a dummy-atom model, or simultaneously fitting both scattering curves to a dummy-residue model, with the imposition of P2 symmetry. The models thus obtained are consistent with the pUL7:pUL51 complex comprising a folded core with an extended region of poorly-ordered amino acids (Fig. 1*C* and 1*D*). In agreement with this, dimensionless Kratky plots of the 1:2 and 2:4 complex SAXS data shows both to have maxima above sR_g_ = √3 (Fig. 1*L*) with extended tails observed in the corresponding probable frequency of real-space distances (p(r) profiles) at longer vector-length distances (Fig 1*K*).

**Figure 1.**
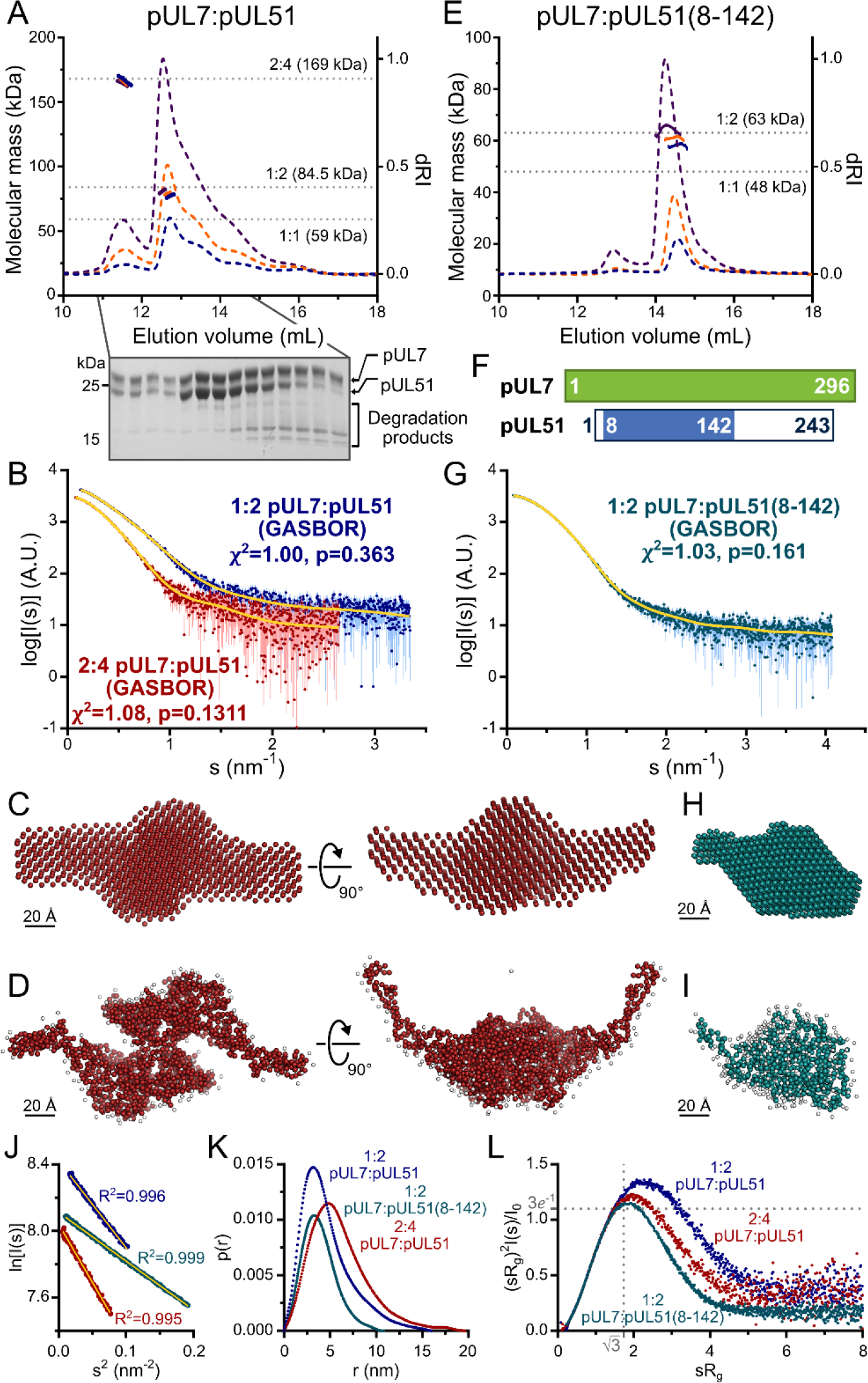
HSV-1 pUL7:pUL51 forms a 1:2 heterotrimeric complex in solution. (A) SEC-MALS analysis of recombinant full-length pUL7:pUL51 complex. Weight-averaged molecular masses (colored solid lines) are shown across the elution profiles (normalized differential refractive index, dRI, colored dashed lines) for samples injected at 2.4, 4.9 and 9.7 mg/mL (blue, orange and purple, respectively). The expected molecular masses for 1:1, 1:2 and 2:4 pUL7:pUL51 complexes are shown as dotted horizontal lines. (B) Averaged SAXS profiles through SEC elution peaks corresponding to 1:2 (blue) and 2:4 (red) complexes of pUL7:pUL51. Fits of representative GASBOR *ab initio* dummy-residue models to the scattering curves for each complex are shown in yellow. χ^2^, fit quality. p, Correlation Map (CorMap) *P*-value of systematic deviations between the model fit and scattering data (46). (C) Refined DAMMIN dummy-atom model reconstruction of the 2:4 pUL7:pUL51 complex, shown in two orthogonal orientations. (D) Representative GASBOR dummy-residue model of the 2:4 pUL7:pUL51 complex, shown in two orthogonal orientations. This model comprises an anti-parallel dimer of heterotrimers, although we note that parallel dimers are also consistent with the scattering data. (E) SEC-MALS of pUL7:pUL51(8-142) complex. Elution profiles and molecular masses are shown as in (A) for recombinant pUL7:pUL51(8–142) injected at 0.6, 1.1 and 3.9 mg/mL (blue, orange and purple, respectively). (F) Schematic representation of pUL7 and pUL51. (G) Averaged SEC-SAXS profile through pUL7:pUL51(8–142) elution peak. Fit of a representative GASBOR *ab initio* dummy-residue model to the scattering curve is shown in yellow. (H) Refined DAMMIN dummy-atom model reconstruction of pUL7:pUL51(8–142) complex. (I) Representative GASBOR dummy-residue model of pUL7:pUL51(8-142). (J) Plot of the Guinier region (sR_g_ < 1.3) for SAXS profiles shown in (B) and (G). The fit to the Guinier equation (yellow) is linear for each curve, as expected for aggregate-free systems. (K) p(r) vs r pofiles for SAXS profiles shown in (B) and (G). (L) Dimensionless Kratky plot of SAXS profiles shown in (B) and (G). The expected maximum of the plot for a compact globular domain that conforms to the Guinier approximation is shown (sR_g_ = √3, (sR_g_)^2^I(s)/I_0_ = 3*e*^−1^, grey dotted lines).

Previous truncation analysis had shown residues 29–170 of pUL51 to be sufficient for pUL7 binding (6). However, neither pUL7 in complex with pUL51 residues 29–170, nor with pUL51 residues 1–170, proved amenable to crystallization. Mass spectrometry analysis identified a smaller protein species, evident whenever the pUL7:pUL51(1–170) was analyzed by SDS-PAGE, as pUL51 residues 8–142. On the assumption that this represented the proteolysis-resistant fragment of pUL51, pUL7 was co-expressed and co-purified with pUL51(8–142). This protein complex could be readily purified and was monodisperse in solution, SEC-MALS showing the pUL7:pUL51(8–142) complex to have a mass of 61.5 ± 3.1 kDa, consistent with a 1:2 complex (calculated mass 63.1 kDa) as observed for full-length pUL7:pUL51 (Fig. 1*E*). SEC-SAXS analysis (Fig. 1*G*) showed the pUL7:pUL51(8–142) complex to be much more compact (R_g_ = 3.0 nm; *D*_*max*_ = 11.5 nm; Fig. 1*K* and Table S1). The Gaussian-like appearances of a dimensionless Kratky plot of the pUL7:pUL51(8–142) scattering data, which is centered on sR_g_ of √3 (Fig. 1*L*), and of the corresponding p(r) profile (Fig. 1*K*) are consistent with the protein having a globular fold. *Ab initio* shape analysis of this data reveals that the pUL7:pUL51(8–142) complex visually resembles the folded core of the full-length complex (Fig. 1*H* and 1*I*).

### Structure of pUL7 in complex with pUL51(8–142)

The pUL7:pUL51(8–142) complex was crystallized and its structure was solved by four-wavelength anomalous dispersion analysis of a mercury acetate derivative. The structure of native pUL7:pUL51(8–142) was refined to 1.83 Å resolution with residuals *R* = 0.195 *R*_free_ = 0.220 and excellent stereochemistry, 99% of residues occupying the most favored region of the Ramachandran plot (Table S2). The asymmetric unit contained four copies of pUL7 residues 11–234 and 253–296 plus eight residues from the C-terminal purification tag (see below) and four copies of pUL51 residues 24–89 and 96–125, the remaining residues of pUL7 and pUL51(8–142) being absent from electron density and presumably disordered.

Strikingly, the molecules of pUL7 and pUL51 in the structure were arranged as a hetero-octamer, with single β-strands from each pUL7 and pUL51 molecule in the asymmetric unit forming a central β-barrel (Fig. 2*A*). Closer inspection revealed that the pUL7 strands in this β-barrel comprised the C-terminal amino acids encoded by the restriction site and from the human rhinovirus 3C protease recognition sequence that remained following proteolytic removal of the GST purification tag. We therefore suspected that this hetero-octameric pUL7:pUL51 arrangement was an artefact of crystallization. SEC-MALS of a pUL7:pUL51(8–142) construct where the purification tag was moved to the N terminus of pUL7, and would thus be unlikely to form the same β-barrel observed in the crystal structure, yielded the same 1:2 pUL7:pUL51 heterotrimeric stoichiometry as observed with C-terminally tagged pUL7 (Fig. S3*A*). Removal of residues 8–40 from pUL51, including residues 24–40 that form part of the β-barrel, yielded a 1:1 heterodimeric complex of pUL7 and pUL51(41–142) as determined by SEC-MALS (Fig. S3*B*), although we note that this truncated complex had reduced solubility. Taken together, these results suggest that pUL7 and pUL51 residues 41–142 assemble to form a heterodimeric ‘core’ complex and that recruitment of the additional pUL51 molecule in the native heterotrimeric complex is mediated by the N-terminal region (residues 8–40) of pUL51.

**Figure 2.**
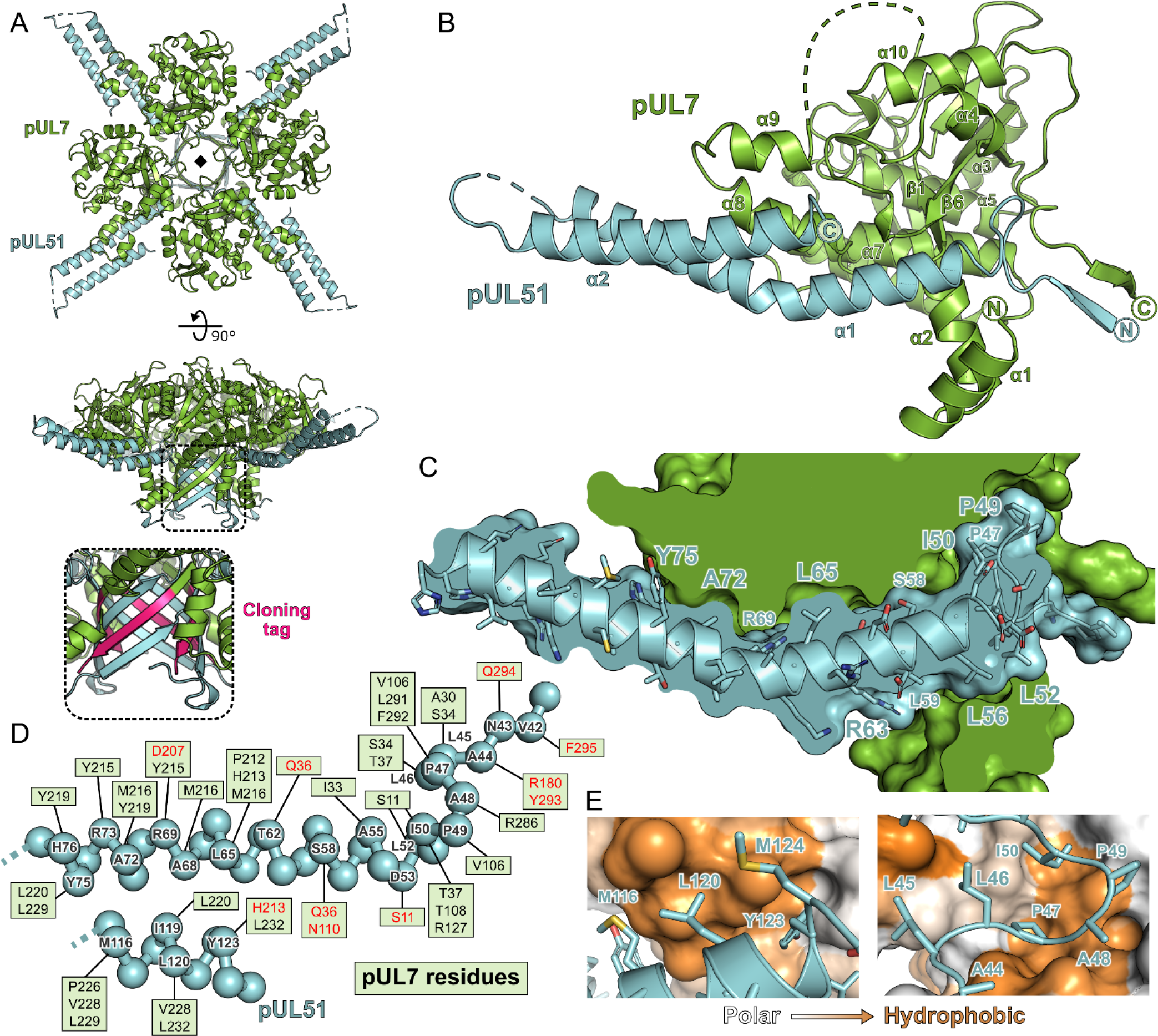
Structure of pUL7 in complex with pUL51. (A) Hetero-octamer of pUL7 and pUL51(8–142) observed in the crystallographic asymmetric unit. pUL7 and pUL51 are shown as green and cyan ribbons, respectively, in two orthogonal orientations. Inset shows residues arising from the pUL7 cloning tag (pink) that form an eight-stranded β-barrel with residues from pUL51. (B) Core heterodimer of pUL7 (residues 11–296) and pUL51 (residues 41–125). Selected secondary structure elements are labelled. (C) ‘Cut-through’ molecular surface representation of pUL7 (green) showing the intimate interaction interface with the hydrophobic loop and helix α1 of pUL51 (cyan). pUL51 side chains are shown as sticks. (D) Molecular interactions between pUL51 (cyan) and pUL7 (boxed residue names). Hydrophobic and hydrogen bond interactions are in black and red typeface, respectively. (E) Molecular surface representation of pUL7, colored by residue hydrophobicity from *white* (polar) to *orange* (hydrophobic). pUL51 is represented as a cyan ribbon with selected side chains shown.

The core heterodimeric complex formed by pUL7 residues 11–296 and pUL51 residues 41–125 is shown in Fig. 2*B*. pUL7 comprises two short N-terminal α-helices followed by a compact globular fold with a mixed α-helical and β-sheet topology containing a central anti-parallel β-sheet and two mostly-buried α-helices that are surrounded by a β-hairpin and additional helices (Fig. S4). Structure-based searches of the protein data bank did not reveal any other domains with a similar fold, which we will henceforth refer to as the Conserved U_L_7(Seven) Tegument Assembly/Release Domain (CUSTARD) fold. pUL51 comprises a hydrophobic loop region followed by a helix-turn-helix. The interaction with pUL7 is extensive and largely hydrophobic in nature (Fig. 2): The hydrophobic loop of pUL51 (residues 45–50, sequence LLPAPI) interacts with pUL7 helix α2 and with a hydrophobic pocket formed by sheets β1 and β6, helices α4 and α7 and the C-terminal tail of pUL7; hydrophobic residues of pUL51 helix α1 interact with a hydrophobic face of pUL7 helix α8; and hydrophobic residues from the C-terminal portion of pUL51 helix α2 interact with hydrophobic residues from pUL7 helices α8 and α9 (Fig. 2 *C*-*E*).

Chemical cross-linking and mass spectrometry was used to further characterize the interaction between pUL7 and pUL51 in solution. As shown in Fig. S5*A*, incubation of the pUL7:pUL51(8–142) complex with either disuccinimidyl sulfoxide (DSSO) or disuccinimidyl dibutyric urea (DSBU) yielded species with masses corresponding to 1:1 or 1:2 pUL7:pUL51 complexes, plus some higher-order species. Analysis of these cross-linked species complexes by MS3 mass spectrometry identified multiple cross-links between pUL7 and pUL51 residues (Table S3). Five of these crosslinks were not compatible with the heterodimer crystal structure, suggesting that they were formed by the other molecule of pUL51 in the heterotrimer, whereas other cross-links could have been formed by either pUL51 molecule. Multiple pseudo-atomic models of the 1:2 pUL7:pUL51(8–142) solution heterotrimer were thus generated by fitting the SAXS profile using the core heterodimer structure, a second copy of pUL51 residues 41–125, and permutations of the feasible chemical cross-linking restraints. Of the 80 models thus generated, half could not simultaneously satisfy all crosslinking restraints and were discarded. The other models all fit the pUL7:pUL51(8–142) SAXS profile well (χ^2^ < 1.26). These models showed the second copy of pUL51 to have the same general orientation relative to pUL7, binding near pUL7 helices α1, α2, α6, α7, and the loop between helices α7 and α8 (Fig. S5*C*). However, the precise orientations of this second pUL51 copy differed, as did the locations of the pUL51 termini. The observed variability is within the resolution limits provided by SAXS, although it may also point to co-existence of alternate conformations, i.e. that the second copy of pUL51 does not adopt one well-defined conformation in solution.

### The interaction between pUL7 and pUL51 is conserved across herpesviruses, but the molecular details of the interface have diverged

The α-, β- and γ-herpesvirus families diverged approximately 400 million years ago (26). Homologues of pUL7 from α-, β- and γ-herpesviruses can be readily identified by their conserved amino acid sequences, despite relatively low sequence identities (HCMV and KSHV homologues share 17% and 16% identity, respectively, with HSV-1 pUL7). The predicted secondary structures of pUL7 homologues from representative α-, β- and γ-herpesviruses that infect humans are very similar to the experimentally-determined secondary structure of HSV-1 pUL7, strongly suggesting that these proteins will adopt the CUSTARD fold (Fig. S2). Similarly, the predicted secondary structures of putative β- and γ-herpesvirus pUL51 homologues closely match the prediction for HSV-1 pUL51 (Fig. S2) despite low sequence identity (HCMV and KSHV homologues sharing 16% and 13% identity, respectively, with HSV-1 pUL51). As the pUL7 and pUL51 homologues conserve secondary structure and, where tested, conserve function in promoting virus assembly, we sought to determine whether the formation of a pUL7:pUL51 complex is conserved across the α-, β- and γ-herpesvirus families.

GFP-tagged pUL7 homologues from human herpesviruses HSV-1, VZV, HCMV or KSHV were co-transfected with mCherry-tagged pUL51 homologues from the same virus into human embryonic kidney (HEK) 293T cells. In all cases, pUL51-mCherry or the relevant homologue could be readily co-precipitated with the GFP-pUL7 homologue, whereas pUL51-mCherry homologues were not efficiently co-precipitated by GFP alone (Fig. 3*A*). The association of pUL7 and pUL51 homologues is therefore conserved across the herpesvirus families.

**Figure 3.**
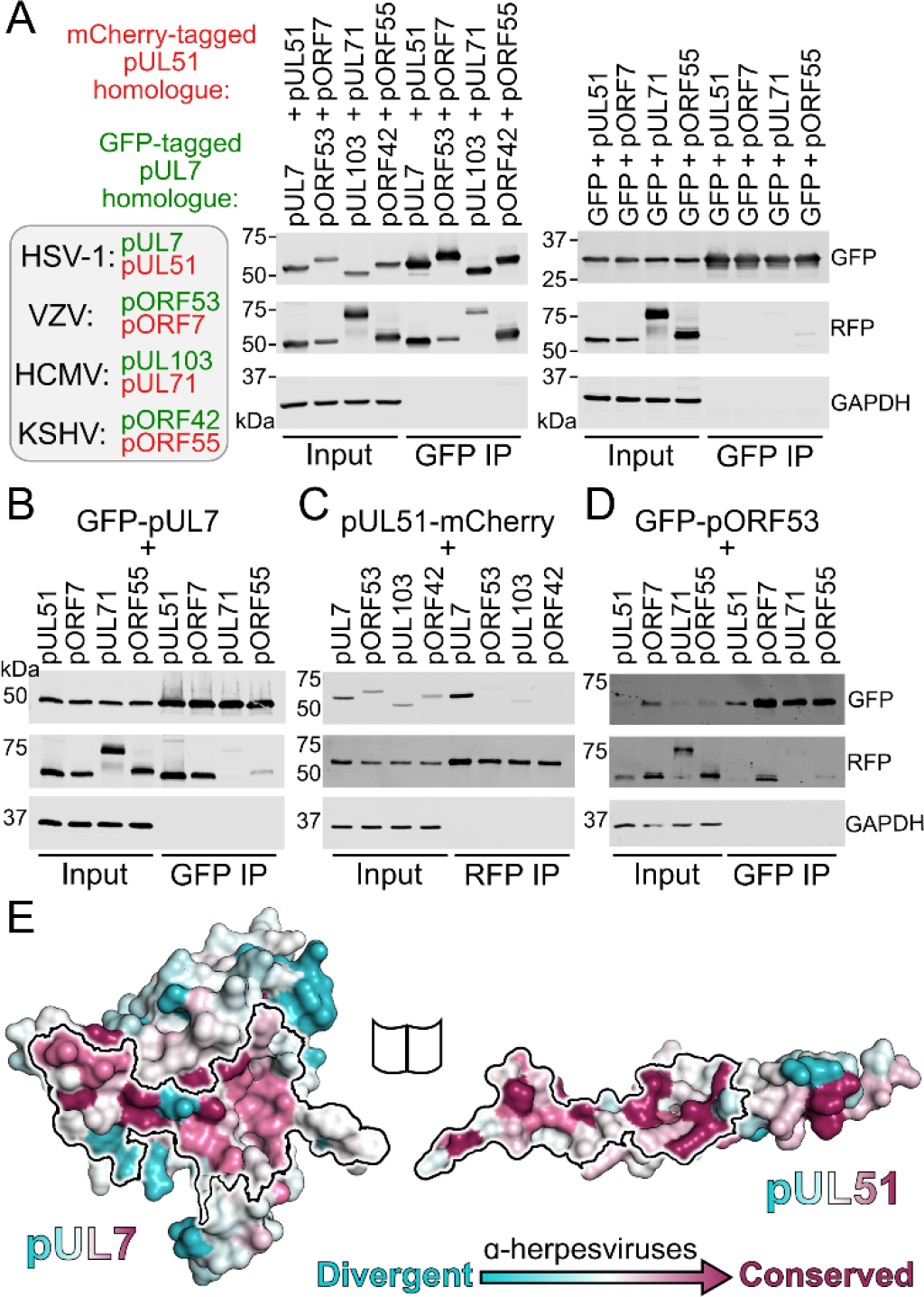
Conservation of the pUL7:pUL51 interaction across herpesviruses. (A-E) HEK 293T cells were co-transfected with GFP-tagged pUL7 homologues from human herpesviruses, or with GFP alone, and with mCherry tagged pUL51 homologues. Cells were lysed 24 h post-transfection and incubated with anti-GFP (A, B, D) or anti-RFP (C, E) resin to capture protein complexes before being subjected to SDS-PAGE and immunoblotting using the antibodies shown. (A) mCherry-tagged homologues of pUL51 are captured by GFP-pUL7 homologues, but not by GFP alone. (B) GFP-pUL7 co-precipitates with pUL51 (HSV-1) and pORF7 (VZV), but not with pUL71 (HCMV) or pORF55 (KSHV). (C) pUL51-mCherry co-precipitates with pUL7 but not with homologues from other herpesviruses. (D) The VZV pUL7 homologue pORF53 co-precipitates with VZV pORF7, but not with pUL51 homologues from other herpesviruses. (E) Molecular surfaces of the pUL7 and pUL51 core heterodimer, colored by residue conservation across the α-herpesviruses. Residues that mediate the pUL7:pUL51 interaction are outlined.

Given the large evolutionary distance between α-, β- and γ-herpesvirus pUL7 and pUL51 homologues, and consequent sequence divergence, it was unclear whether the molecular details of the interaction between these proteins would be conserved. GFP-tagged pUL7 was thus co-transfected with mCherry-tagged pUL51 from HSV-1 or with mCherry-tagged homologues from VZV, HCMV or KSHV. Co-precipitation was observed for HSV-1 pUL51 and for pORF7 from VZV, an α-herpesvirus, but not for the homologues from HCMV or KSHV (Fig. 3*B*). This suggested that the pUL7:pUL51 molecular interface is partially conserved within the α-herpesvirus family, but not across families. VZV pORF53 and pORF7 share 33% and 35% identity with HSV-1 pUL7 and pUL51, respectively. Mapping the conservation of α-herpesvirus pUL7 sequences onto the HSV-1 pUL7 structure reveals several regions of conservation that overlap with the binding footprint in pUL51 in the core heterodimeric complex (Fig. 3*E*). However, capture of pUL51-mCherry did not result in co-precipitation of the VZV pUL7 homologue pORF53, nor did capture of GFP-pORF53 result in co-precipitation of HSV-1 pUL51 (Fig. 3 *C* and *D*). We therefore conclude that, while the pUL7:pUL51 interface is partially conserved across α-herpesviruses, there has been co-evolution of pUL7 and pUL51 homologues such that the interaction interfaces are distinct at a molecular level.

To test whether the core heterodimeric pUL7:pUL51 interaction interface is subject to co-evolutionary change, a matrix of 63 interacting pairs of residues (one from each protein) was generated by manual inspection of the binding interface. The amino acids carried at these sites across an alignment of pUL7 and pUL51 homologues from 199 strains of α-herpesvirus were tested for correlated changes. Initially, 35 of the 63 interacting-residue pairs where homology could be confidently assigned were analyzed, results being compared to a null distribution determined from 10^6^ data sets where interacting sites were paired at random. True pairings showed more correlated change than 94% of the randomized pairings and the results were little changed when different subsets of the data, including fewer strains and more interactions, were analyzed (Table S4). This is suggestive evidence for co-evolution of the interaction interface across the α-herpesviruses. Similar analysis was attempted to probe for co-evolution of the core pUL7:pUL51 interaction interface across all herpesviruses, but the extensive sequence divergence confounded the confident assignment of interacting amino acid pairs (only 12 pairs could be confidently assigned) and so the subsequent analysis was underpowered.

In addition to interacting with pUL7, it has previously been shown that HSV-1 pUL51 is able to interact with the protein pUL14 (25) and that mutation of pUL51 amino acids Ile111, Leu119 and Tyr123 to alanine disrupts this interaction. Residues Leu119 and Tyr123 are completely buried in the interface between pUL7:pUL51 in the core heterodimer structure, interacting with residues from pUL51 helix α2 and from pUL7 helices α8 and α9 (Fig. S6*A*). Such burial would preclude simultaneous binding of these residues to pUL7 and pUL14. However, the second copy of pUL51 in the solution heterotrimer may be capable of binding pUL14, or pUL14 may compete with pUL7 for binding to pUL51. To test these hypotheses, pUL51-mCherry was co-transfected into mammalian cells together with GFP-pUL7 and/or myc-pUL14 and then captured using mCherry affinity resin. While GFP-pUL7 was readily co-precipitated, we could not detect co-precipitation of myc-pUL14 with pUL51-mCherry either in the presence or absence of GFP-pUL7 (Fig. S6*B*).

### Association of pUL7:pUL51 homologues to trans-Golgi membranes is conserved but association with focal adhesions is not

In addition to the roles of the pUL7 and pUL51 in promoting virus assembly, which appear to be conserved across herpesviruses, the HSV-1 pUL7:pUL51 complex has been shown to interact with focal adhesions to stabilize the attachment of cultured cells to their substrate during infection (6). To probe whether focal adhesion binding is a conserved property of pUL7:pUL51 homologues, GFP-tagged pUL7 and mCherry-tagged pUL51 (or homologous complexes) were co-transfected into HeLa cells. As previously observed, HSV-1 pUL7:pUL51 complex co-localizes with both TGN46, a *trans*-Golgi marker, and with paxillin and zyxin at the cell periphery, markers of focal adhesions (Fig. 4, S7, S8). VZV pORF53:pORF7, HCMV pUL103:pUL71 and KSHV pORF42:pORF55 all co-localize with TGN46 at *trans*-Golgi membranes (Fig. 4). However, these homologues do not co-localize with paxillin or zyxin at focal adhesions (Fig. S7 and S8).

**Figure 4.**
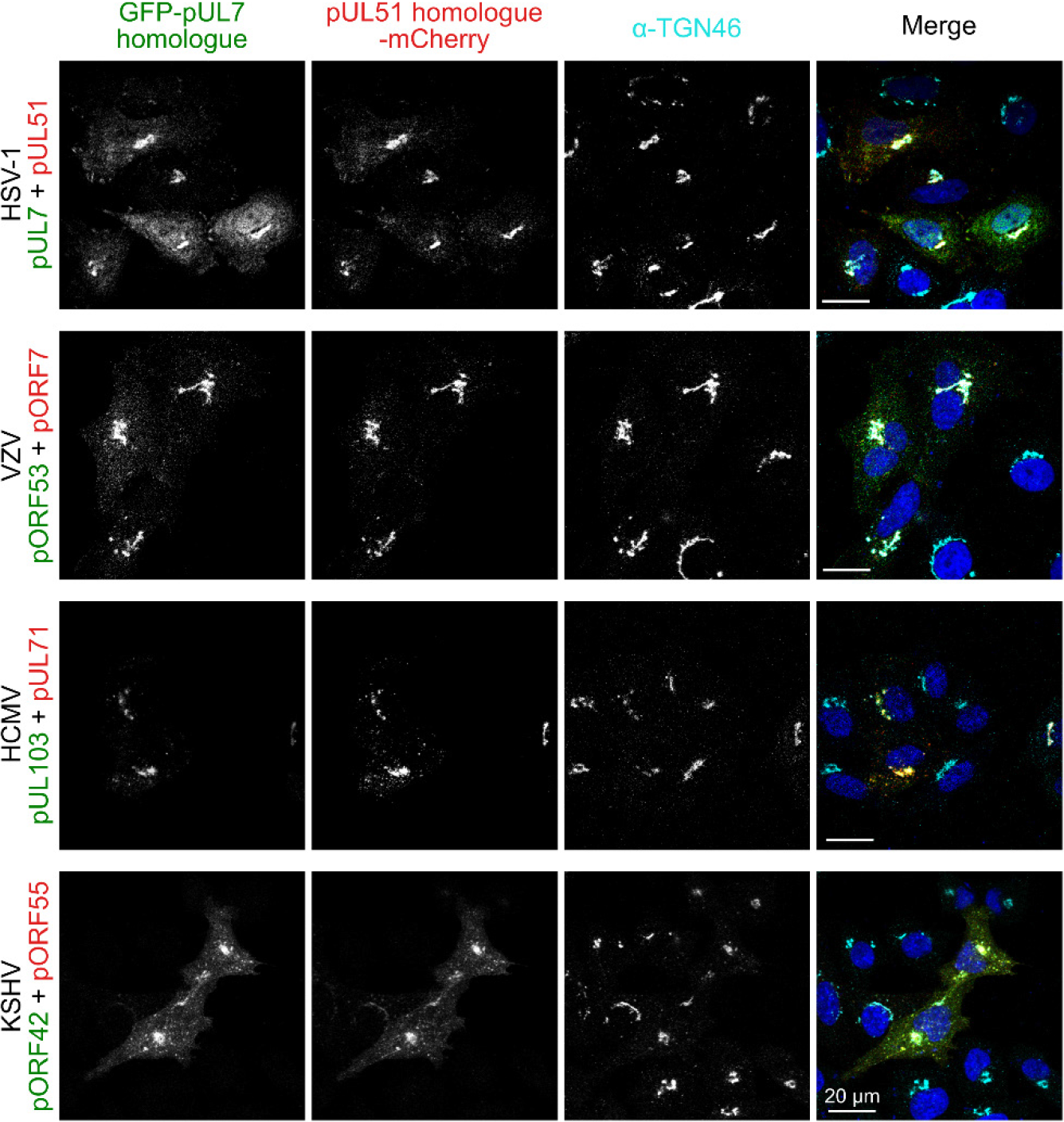
Co-localization of the pUL7:pUL51 complex with *trans*-Golgi membranes is conserved across human herpesviruses. HeLa cells were co-transfected with GFP-pUL7 and pUL51-mCherry, or with similarly-tagged homologues from VZV, HCMV and KSHV. Cells were fixed 24 hours post transfection and immunostained using the *trans*-Golgi marker protein TGN46 before imaging by confocal microscopy. Co-localization between the GFP, mCherry and far-red (TGN) fluorescence is observed in cells transfected with either HSV-1 pUL7:pUL51 or with the homologous complexes from VZV, HCMV and KSHV. HSV-1 pUL7 and pUL51 also co-localize with striated cell peripheral structures (focal adhesions, see Fig. S7 and S8).

## Discussion

We present here the structure of HSV-1 pUL7 in complex with pUL51. In solution this complex forms a 1:2 heterotrimer that is capable of forming higher-order oligomers (Fig. 1). The C-terminal region of pUL51 is predicted to be disordered (Fig. S2) and is extended in solution (Fig. 1 *C*, *D* and *L*), consistent with this region having little intrinsic structure. The crystal structure of pUL7 in complex with pUL51(8–142) shows pUL7 to comprise a single compact globular domain that adopts a previously-unobserved CUSTARD fold (Fig. 2 and Fig. S4). A single molecule of pUL51 is bound to pUL7 in this crystal structure via an extended hydrophobic interface that is largely conserved across α-herpesviruses (Fig. 3) and there is evidence that residues at the interface are co-evolving (Table S4). Most of the pUL7-interacting residues lie within the hydrophobic loop and helix α1 of pUL51 (residues 45–88), consistent with a recent report that pUL51 residues 30–90 are sufficient for the interaction with pUL7 in transfected cells (27). Recruitment of the second copy of pUL51 to the pUL7:pUL51 complex in solution requires pUL51 residues 8–40 (Fig. 1 and S3), consistent with observations that the equivalent N-terminal region of the HCMV pUL51 homologue pUL71 is required for its self-association both *in vitro* and in cultured cells (28), and that VZV pUL51 homologue pORF7 can also form higher-order oligomers (12).

The interaction between pUL7 and pUL51 homologues is conserved across all three families of herpesvirus (Fig. 3*A*), as is the association of these complexes with *trans*-Golgi compartments in cultured cells (Fig. 4), but of the complexes tested only HSV-1 pUL7:pUL51 associates with focal adhesions in cultured cells (Fig. S7 and S8). The conserved association of pUL7:pUL51 complexes with *trans*-Golgi membranes is consistent with a conserved role for this complex in herpesvirus assembly. Assembly of HSV-1 occurs at juxtanuclear membranes that contain cellular trans-*Golgi* and endosomal marker proteins (4, 29) and that are derived, at least in part, from recycling endosomes (30). Similarly, HCMV assembly occurs at viral assembly compartments that contain *trans*-Golgi marker proteins (5, 31) and mutation of the pUL71 Yxxϕ motif, which mediates recycling from the plasma membrane via recognition by AP2 (32), causes re-localization of pUL71 to the plasma membrane and prevents efficient HCMV assembly (18). Given the conservation of the pUL7:pUL51 interaction, the conserved localization of this complex to *trans*-Golgi membranes, and the established evidence supporting roles for pUL7 or pUL51 homologues in virus assembly (6, 9, 13, 14, 19–21, 23, 24), we propose that pUL7 and pUL51 form a complex that is conserved across herpesviruses and functions to promote virus assembly by stimulating cytoplasmic envelopment of nascent virions.

In addition to binding pUL7, pUL51 has been shown to interact with pUL14 and disruption of this interaction inhibits virus assembly (25). We did not observe this interaction in co-precipitation experiments performed using transfected cells in the absence of infection (Fig S6*B*). Oda *et al.* (25) reported that mutations of pUL51 residues Ile111, Leu119 and Tyr123 to alanine disrupted an interaction between pUL51 and pUL14. However, Leu119 and Tyr123 are completely buried in the pUL7:pUL51 interaction interface (Fig. S6*A*) and such mutation would be likely to severely destabilize this interaction. As all the experiments of Oda and colleagues (25) were performed using infected cells or infected-cell lysates, it seems likely that the observed interaction between pUL51 and pUL14 is not direct but is instead mediated by other herpesvirus proteins, and that such an interaction may require binding of pUL51 to pUL7.

While the pUL7 CUSTARD fold has not been observed previously, frustrating attempts to infer function by analogy, the helix-turn-helix fold of pUL51 residues 41–125 is a common feature of many proteins. Of the proteins identified by structure-based searches, the similarity to human CHMP4B, a component of the ESCRT-III membrane-remodeling machinery, is particularly notable given the role of pUL51 and homologues in stimulating virus wrapping (6, 13, 14, 20). CHMP4B and homologues are required for the efficient fusion of membrane leaflets during vesicle budding into organelle lumens, cytokinetic abscission, nuclear envelope closure, and budding of some enveloped viruses (33). Helices α1 and α2 of pUL51 superpose onto human CHMP4B (34) with 1.2 Å root-mean-squared deviation across 59 C^α^ atoms (Fig. 5*A*). pUL51 also resembles the structures of yeast and fly CHMP4B homologues Snf7 (35) and Shrub (36) and pUL51 can be superposed onto either structure with 1.5 Å root-mean-squared deviation across 57 C^α^ atoms (Fig. 5*B* and *C*). In these Snf7 and Shrub structures the conserved helix α3 of the ESCRT-III core domain is continuous with helix α2 and is longer than pUL51 helix α2. However, we note that the region of pUL51 immediately following helix α2 is predicted to be helical (Fig. S2) and in ESCRT-III structures helix α3 is known to be mobile, adopting both ‘closed’ and ‘open’ (extended) conformations (33). The polymerization of CHMP4B is known to be regulated by association with CC2D1A in humans (34) and Lgd in flies (37). Superposition of the pUL7:pUL51 core heterodimer onto Shrub shows that pUL7 occupies the space that would be occupied by the adjacent Shrub molecule of a putative Shrub homopolymer (Fig. 5*D*) (36). Similarly, the DM14-3 domain of Lgd, which is sufficient to bind Shrub *in vitro* and prevents Shrub polymerization (37), occupies a similar space to helices α8 and α9 of pUL7 (Fig 5*E*). It is therefore notable that pUL51 is prone to self-association when expressed in the absence of pUL7 (Fig. S1).

**Figure 5.**
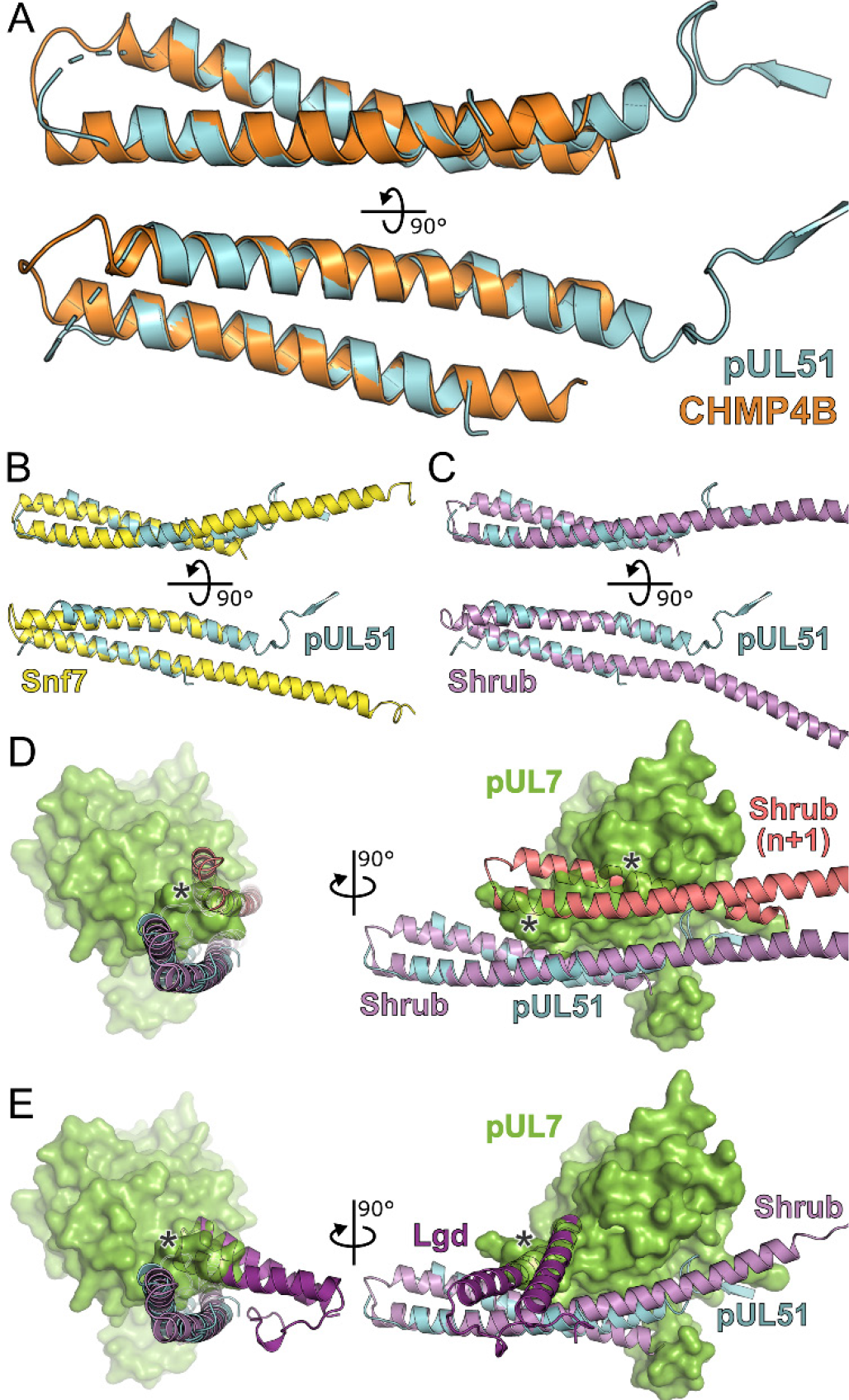
Structural similarity of HSV-1 pUL51 to cellular ESCRT-III proteins. (A) pUL51 (cyan) is shown superposed on helical hairpin (conserved helices α1 and α2) of human CHMP4B (orange; PDB 4ABM) (34). (B and C) pUL51 (cyan) superposed on conserved helices α1 and α2 of CHMP4B homologues (B) yeast Snf7 (yellow; PDB 5FD7) (35) and (C) fly Shrub (violet; PDB 5J45) (36). (D and E) The pUL7:pUL51 core heterodimer is shown superposed onto (D) two subunits of the putative Shrub homopolymer (violet and pink; PDB ID 5J45) (36), or (E) the complex of Shrub with the regulatory DM14-3 domain of Lgd (purple; PDB 5VO5) (37). pUL7 is shown as a green molecular surface. Spatial overlap between pUL7 and (D) the second subunit of Shrub, or (E) the Lgd DM14-3 domain, is denoted by asterisks.

Activity of ESCRT-III components and VPS4, the AAA-ATPase that promotes fusion and dissociates ESCRT-III (33, 38), are known to be required for efficient assembly of HSV-1 (39, 40). However, a recent study of HCMV identified that neither VPS4 nor mammalian ESCRT-III components CHMP4B or CHMP6 are required for assembly, and the authors hypothesized that pUL71 may act as a viral ESCRT-III homologue (41). The structural similarity between pUL51 and CHMP4B supports this hypothesis: The predicted secondary structure of pUL51 is conserved across herpesviruses, as is the interaction with pUL7, suggesting that the N-terminal regions of all pUL51 homologues will conserve this CHMP4B-like fold. This hypothesis is further supported by the recent observation that mutation of four lysine residues near the C terminus of pUL71 results in the accumulation of HCMV particles in membrane buds with narrow necks (42), reminiscent of the stalled budding profiles observed for HIV-1 when ESCRT-III activity is perturbed (43) or when HSV-1 infects cells expressing a dominant negative form of VPS4 (39). The mechanism by which herpesviruses recruit ESCRT-III to tegument-wrapped capsids in order to catalyze cytoplasmic envelopment remains unknown (44). Given the structural and functional homology between pUL51 and ESCRT-III components, it is tempting to speculate that pUL51 homologues act as viral ESCRT-III components, either alone (HCMV) or in concert with cellular proteins like CHMP4B (HSV-1), and that pUL7 homologues may act like CC2D1A to regulate pUL51 homopolymerisation in infected cells.

## Materials and Methods

Full materials and methods are provided in *SI materials and methods*. Briefly, proteins pUL7 and pUL51 (with residue Cys9 substituted for serine) from HSV-1 strain KOS, or truncations thereof, were co-expressed in *Escherichia coli* and co-purified by affinity capture and SEC. MALS data were acquired inline following SEC using an S200 increase 10/300 column (GE Healthcare) equilibrated in 20 mM tris pH 7.5, 0.2 M NaCl, 3% (v/v) glycerol, 0.25 mM tris(2-carboxyethyl)phosphine. SEC-SAXS data were acquired at EMBL-P12 bioSAXS beam line (45) and analyzed using the ATSAS package (46). *Ab initio* bead models of pUL7:pUL51(full-length or 8–142) were generated using GASBOR (47) and DAMMIN (48), and pseudo-atomic models of pUL7:pUL51(8–142) were generated using CORAL (46). Crosslinking was performed by incubating purified pUL7:pUL51(8–142) with DSSO or DSBU and mass spectra were acquired on an Orbitrap Fusion Lumos (Thermo Fisher). pUL7:pUL51(8–142) was crystallized in sitting or hanging drops by mixing 1 μL of 5.3 mg/mL protein with 0.5 μL of 0.5 M benzamidine hydrochloride and 1 μL of reservoir solution containing 0.15 mM sodium citrate pH 5.5, 12% (v/v) 2-methyl-2,4-pentanediol, 0.1 M NaCl and equilibrating against 200 μL reservoirs at 16°C for at least one week. Diffraction data were processed using DIALS (49) and AIMLESS (50), the structure was solved by four-wavelength anomalous dispersion using a mercury derivative with CRANK2 (51), and was refined against the high-resolution native diffraction data using autoBUSTER (52). HEK 293T cells were transfected with GFP- and mCherry tagged pUL7 and pUL51, or homologues from other herpesviruses, which were subsequently precipitated using GFP-Trap or RFP-Trap affinity resins (Chromotek) and visualized by immunoblotting. HeLa cells co-transfected with GFP- and mCherry tagged pUL7 and pUL51, or homologues from other herpesviruses, were fixed and imaged by confocal microscopy following staining for markers of the *trans*-Golgi network (TGN46) or focal adhesions (zyxin or paxillin).

## Data availability

Crystallographic coordinates and structure factors have been deposited in the Protein Data Bank, www.pdb.org [PDB ID code 6T5A], and raw diffraction images have been deposited in the University of Cambridge Apollo repository (https://doi.org/10.17863/CAM.44914). SAXS data, *ab initio* models and pseudo-atomic models have been deposited into the Small-Angle Scattering Biological Data Bank (SASBDB) (53) under the accession codes SASDG37 (pUL7:pUL51(8–142) heterotrimer), SASDG47 (pUL7:pUL51 heterohexamer) and SASDG57 (pUL7:pUL51 heterotrimer). Other materials will be provided upon request.

## Acknowledgments

HCMV and KSHV cDNA were kind gifts of John Sinclair and Mike Gill. We thank Janet Deane for access to MALS equipment, Janet Deane, Chris Hill and the mentors at the DLS-CCP4 Data Collection and Structure Solution Workshop 2018 for helpful discussions, Len Packman for peptide fingerprinting mass spectroscopy analysis, Susanna Colaco for superb technical assistance, Diamond Light Source for access to beamlines I03 and I04 under proposal mx15916, and EMBL for access to the bioSAXS beamline P12 under proposal HH-SAXS-911. Remote synchrotron access was supported in part by the EU FP7 infrastructure grant BIOSTRUCT-X (Contract No. 283570) and access to P12 was supported by iNEXT funded by the Horizon 2020 programme of the European Commission (grant number 653706). A Titan V graphics card used for this research was donated by the NVIDIA Corporation. BGB is a Wellcome Trust PhD student, DJO was supported by a John Lucas Walker Studentship, and MFA was supported by Commonwealth Scholarship Commission PhD scholarship (BDCA-2014-7). This work was supported by a Sir Henry Dale Fellowship (098406/Z/12/B), jointly funded by the Wellcome Trust and the Royal Society (to SCG).

