## Supplemental material for "Structure of herpes simplex virus pUL7:pUL51, a conserved complex required for efficient herpesvirus assembly"

### SI Materials and Methods

*Protein production*

Herpes simplex virus (HSV)-1 strain KOS protein pUL51 (UniProt ID D3YPL0), either with the wild-type sequence or where residue Cys9 (the palmitoyl group acceptor) had been substituted with serine, was expressed with an N-terminal MetAlaHis$_6$ tag and purified by $Ni^{2+}$ affinity capture and size-exclusion chromatography as described in (1). The complex of HSV-1 strain KOS proteins pUL7 (UniProt ID A0A110B4Q7) and pUL51, or truncations thereof, were co-expressed in the *Escherichia coli* strain T7 express *lysY/I$^q$* (New England BioLabs) using the polycistronic vector pOPC (2). The nucleotide sequence of pUL7 had been optimized to enhance recombinant expression (GeneArt) and, where present, residue Cys9 of pUL51 had been substituted with serine. For all experiments except Fig. S3, pUL7 was fused to a C-terminal human rhinovirus 3C protease recognition sequence and GST purification tag. For Fig. S3*A*, the GST and 3C recognition sequence were fused to the N terminus of pUL7. Bacteria were cultured in 2×TY medium, recombinant proteins being expressed overnight at 22°C following addition of 0.4 mM isopropyl β-D-1-thiogalactopyranoside.

Bacterial cell pellets were resuspended in lysis buffer (50 mM sodium phosphate pH 7.5, 500 mM NaCl, 0.5 mM $MgCl_2$, 1.4 mM β-mercaptoethanol, 0.05% Tween-20) supplemented with 400 U bovine pancreas DNase I (Merck) and 200 µL EDTA-free protease inhibitor cocktail (Merck) at 4°C. Cells were lysed using a TS series cell disruptor (Constant Systems) at 24 kPSI and the lysate was cleared by centrifugation at 40,000×g for 30 min at 4°C. Cleared lysate was incubated with glutathione sepharose 4B resin (GE Healthcare) equilibrated in GST wash buffer (50 mM sodium phosphate pH 7.5, 300 mM NaCl, 1 mM dithiothreitol (DTT)) for 1 h at 4°C before being applied to a column and washed with >10 column volumes (c.v.) of GST wash buffer. To remove contaminating nucleic acids, pUL7:pUL51 complexes were resuspended in 25 mM sodium phosphate pH 7.5, 150 mM NaCl, 0.5 mM DTT, 1 mM $MgCl_2$ and incubated with 2000 U of benzonase (Merck) for 1 h at room temperature before being applied to a column, washed with 8 c.v. of 50 mM sodium phosphate pH 7.5, 1M NaCl, and then washed with 4 c.v. of GST wash buffer. Bound protein was eluted using GST wash buffer supplemented with 25 mM reduced L-glutathione, concentrated, and further purified by size-exclusion chromatography (SEC) using an S200 16/600 column (GE Healthcare) equilibrated in 20 mM tris pH 7.5, 200 mM NaCl, 1 mM DTT. The GST tag was removed by supplementing the pooled SEC fractions containing pUL7:pUL51 complex with 0.5 mM EDTA and then incubating with 40 µg of GST-tagged human rhinovirus 3C protease. Free GST and uncleaved GST-tagged pUL7 were captured using glutathione sepharose resin and the cleaved complex was again subjected to SEC using S200 16/600 or 10/300 columns (GE Healthcare) equilibrated in 20 mM tris pH 7.5, 200 mM NaCl, 1 mM DTT, 3% (v/v) glycerol. Purified pUL7-pUL51 was concentrated, plunge-frozen in liquid nitrogen as small (<100 µL) aliquots, and stored at -80°C. Protein concentrations were estimated from absorbance at 280 nm using calculated extinction coefficients where pUL7 and pUL51 were assumed to be present in 1:2 molar ratios for all complexes except for pUL7:pUL51(41-142), where an equimolar ratio was assumed.

*Multi-angle light scattering*

Multi-angle light scattering (MALS) experiments were performed immediately following SEC (SEC-MALS) by inline measurement of static light scattering (DAWN 8+; Wyatt Technology), differential refractive index (Optilab T-rEX; Wyatt Technology), and UV absorbance (1260 UV; Agilent Technologies). Samples (100 µL) were injected onto an S200 Increase column (GE Healthcare) equilibrated in SEC buffer (with 1 mM tris(2-carboxyethyl)phosphine (TCEP) substituted for DTT) at 0.4 mL/min. Molecular masses were calculated using ASTRA 6 (Wyatt Technology) and figures were prepared using Prism 7 (GraphPad).

*Small-angle X-ray scattering and* ab initio *modelling*

Continuous flow small-angle X-ray scattering (SAXS) experiments were performed immediately following SEC with in-line MALS and dynamic light scattering (SEC-SAXS-MALS-DLS), at EMBL-P12 bioSAXS beam line (PETRAIII, DESY, Hamburg) (3, 4). Scattering data (I(s) versus s, where $s = 4\pi \sin\theta/\lambda$ $nm^{-1}$, 2θ is the scattering angle, and λ is the X-ray wavelength, 0.124 nm) were recorded using a Pilatus 6M detector (Dectris) with 1 s sample exposure times for a total of 3,600 data frames spanning the entire course of the SEC separation. 90 µL of purified pUL7:pUL51 (8 mg/mL) or pUL7:pUL51(8–142) (4.5 mg/mL) was injected at 0.5 mL/min onto an S200 Increase 10/300 column (GE Healthcare) equilibrated in 20 mM 4-(2-hydroxyethyl)-1-piperazineethanesulfonic acid (HEPES) pH 7.5, 200 mM NaCl, 3% (v/v) glycerol, 1 mM DTT (pUL7:pUL51) or 20 mM tris pH 7.5, 200 mM NaCl, 3% (v/v) glycerol, 0.25 mM TCEP (pUL7:pUL51(8–142)). SAXS data for the pUL7:pUL51 complex, which eluted as two peaks, were recorded from macromolecule-containing and -free fractions as follows: heterohexamers (frames 1236–1283 s), heterotrimers (frames 1382–1416 s) and solvent blank (spanning pre- and post-sample elution frames). For the pUL7:pUL51(8–142) complex, the SEC-SAXS experiment was performed three times as follows: heterotrimers (frames 1779–1886 s, 1785–1876 s and 1781–1880 s for the three experiments, respectively) and solvent blanks (spanning pre- and post-sample elution frames). Primary data reduction was performed using CHROMIXS (5) and 2D-to-1D radial averaging was performed using the SASFLOW pipeline (6). Buffer frames were tested for statistical equivalence using all pairwise comparison CorMap p values set at a significance threshold (α) of 0.01 (7) before being averaged to generate a final buffer scattering profile and subtracted from the relevant macromolecule elution peaks. Subtracted data blocks producing a consistent $R_g$ through the elution profile (as evaluated using the Guinier approximation (8)) were scaled and checked for similarity using CorMap before being averaged to produce the final reduced 1D scattering profiles. For the pUL7:pUL51(8–142) construct, the averaged scattering profiles obtained for the three repeated SEC-SAXS measurements underwent additional scaling and final combined averaging. Primary data processing, including all CorMap calculations, was performed in PrimusQT of the ATSAS package (9). Molecular weight estimates were calculated using the datporod (Porod volume) (9), datmow (10), datvc (11) and Bayesian consensus modules (12) of the ATSAS package. Indirect inverse Fourier transform of the SAXS data and the corresponding probable real space-scattering pair distance distributions (p(r) versus r profile) were calculated using GNOM (13), from which the $R_g$ and $D_{max}$ were determined. In addition, the *a priori* assessment of the non-uniqueness of scattering data was performed using AMBIMETER (14). SAXS data collection and analysis parameters are summarized in Table S1. *Ab initio* modeling was performed using GASBOR (15) and DAMMIN (16). For pUL7:pUL51, reciprocal space intensity fitting accounting for oligomeric equilibrium with P2 symmetry imposed (GASBORMX) was used to simultaneously fit the 1:2 (heterotrimer) and 2:4 (heterohexamer) pUL7:pUL51 SAXS profiles. The two SEC-elution peaks contained heterohexamer:heterotrimer volume fractions of 1.0:0.0 and 0.2:0.8, respectively, as determined by GASBORMX. Because SAXS data can be ambiguous with respect to shape restoration, DAMMIN and GASBOR were run 20 times and the consistency of the individual models was evaluated using the normalized spatial discrepancy (NSD) metric (17). Dummy-atom models were clustered using DAMCLUST (17), averaged using DAMCLUST (pUL7:pUL51) or DAMAVER (pUL7:pUL51(8–142)), and refined using DAMMIN. For the pUL7:pUL51 heterohexamer three clusters were identified, which visually corresponded to parallel or anti-parallel dimers of heterotrimers, whereas for the pUL7:pUL51(8–142) heterotrimer all models formed a single cluster. The refined dummy-atom models that best fit the SAXS profile (lowest $\chi^2$) are shown in Fig. 1.

*Cross-linking and mass spectrometry*

Purified pUL7:pUL51(8–142) at 1 mg/mL (16.4 µM) in SAXS buffer was incubated with 20- to 100-fold molar excess of disuccinimidyl sulfoxide (DSSO; ThermoFisher) or disuccinimidyl dibutyric urea (DSBU; ThermoFisher) dissolved in DMSO, or with DMSO carrier alone (the final DMSO concentration remaining below 2% (v/v) in all cases). Reactions were incubated at room temperature for 30 min before quenching by addition of 1 M tris pH 7.5 to a final tris concentration of 20 mM. Samples were separated by SDS-PAGE using a 4–12% Bolt Bis-Tris gel (ThermoFisher) in MOPS running buffer and stained with InstantBlue Coomassie Protein Stain (Expedeon) according to the manufacturers' instructions. Cross-linked samples corresponding to pUL7:pUL51(8–142) heterodimers (1:1) or heterotrimers (1:2) were excised, reduced, alkylated and digested in-gel using trypsin. The resulting peptides were analyzed using an Orbitrap Fusion Lumos coupled to an Ultimate 3000 RSLC nano UHPLC equipped with a 100 µm ID × 2 cm Acclaim PepMap Precolumn and a 75 µm ID × 50 cm, 2 µm particle Acclaim PepMap RSLC analytical column (ThermoFisher Scientific). Loading solvent was 0.1% formic acid (FA) with analytical solvents A: 0.1% FA and B: 80% (v/v) acetonitrile (MeCN) + 0.1% FA. Samples were loaded at 5 µL/min, loading solvent for 5 minutes before beginning the analytical gradient. The analytical gradient was 3% to 40% B over 42 minutes, rising to 95% B by 45 minutes, followed by a 4-minute wash at 95% B, and finally equilibration at 3% solvent B for 10 minutes. Columns were held at 40ºC. Data was acquired in a DDA fashion with MS3 triggered by a targeted mass difference. MS1 was acquired from 375 to1500 Th at 60,000 resolution, $4\times10^5$ AGC target and 50 ms maximum injection time. MS2 used quadrupole isolation at an isolation width of m/z 1.6 and CID fragmentation (25% NCE). Fragment ions were scanned in the Orbitrap with $5\times10^4$ AGC target and 100 ms maximum injection time. MS3 was triggered by a targeted mass difference of 25.979 for DSBU and 31.9721 for DSSO with HCD fragmentation (30% NCE) and fragment ions scanned in the ion trap with an AGC target of $2.0 \times 10^4$.

Raw files were process using XLinkX 2.2 in Proteome Discoverer 2.2.0.388 (ThermoFisher). MS2 or MS3 spectra were selected based on the identification of either DSSO (K +158.004 Da) or DSBU (K +196.085 Da) and then processed in two workflows in parallel with the following parameters. Workflow 1: XlinkX Search against a database containing an HSV-1 proteome (downloaded 14.14.2016), *E. coli* proteome (downloaded 06.09.2019 with OPGE removed) and 246 common contaminants; full trypsin digestion; carbamidomethyl static modification of cysteines; oxidation variable modification of methionines; 1% FDR using XlinkX validator Percolator. Workflow 2: spectra filtered for either MS2 or MS3 scans with each set searched separately using Mascot against a database containing an *E. coli* proteome (downloaded 06.09.2019 with OPGE removed) with 246 common contaminants, and HSV-1 proteome (downloaded 14.14.2016); PSM validator Max. Delta Cn = 0.05. Statistical validation of identified cross-link peptides from both workflows was carried out by a joint consensus workflow.

*Pseudo-atomic modelling of pUL7:pUL51(8–142) heterotrimer SAXS profile*

Pseudo-atomic modelling of the pUL7:pUL51(8–142) heterotrimer was performed using CORAL (9). The core heterodimer structure, comprising pUL7 and pUL51(41–125), was fixed in this model and a second copy of pUL51(41–125) was free to move. To include *a priori* information about predicted secondary structure (Fig. S2), the pUL51(8–142) sequence was modelled by I-TASSER (18) using pUL51 residues 41–125 from core heterodimer structure as a template. Secondary structural (helical) elements from the I-TASSER model were included for regions of pUL51 that were disordered in the crystal structure (residues 8–23 and 126–142) or involved in the artefactual interaction with the pUL7 purification tag (residues 24–40). DSSO and DSBU cross-links were used to generate maximal inter-residue distance restraints of 26.1 and 28.3 Å, respectively (19). Cross-links between residues of pUL7 and pUL51 that are not feasible based on the core heterodimer structure were assumed to be between pUL7 and the additional copy of pUL51. Cross-links that could not be assigned unambiguously (e.g. cross-links between pUL51 residues that could be either inter- or intra-molecular) were permuted and all possible restraint geometries were tested by modelling against the pUL7:pUL51(8–142) SAXS profile ($s < 3.2$ nm$^{-1}$). The final distribution of target function (F) values was clearly bimodal: models from the cluster with higher F values were unable to simultaneously satisfy the provided cross-link restraints and the SAXS data, and were thus discarded. Remaining models were assessed for fit to the SAXS profile ($\chi^2$) using CRYSOL.

*X-ray crystallography*

Crystals of pUL7:pUL51(8–142) were cryoprotected by brief immersion in reservoir solution supplemented with 20% (v/v) glycerol before flash cryo-cooling by plunging into liquid nitrogen. For multiple-wavelength anomalous dispersion (MAD) phasing experiments, 1 µL of 1 mM mercury(II) acetate in reservoir solution was added to the mother liquor and incubated at 16°C for 4 h before cryoprotection and cryo-cooling as described above. Diffraction data were recorded at 100 K on a Pilatus3 6M detector (Dectris) at Diamond Light Source beamline I03. Images were processed using DIALS (20), either using the DUI graphical interface (21) for the native dataset or the xia2 automated processing pipeline (22) for the mercury derivative datasets. Scaling and merging was performed using AIMLESS (23) and data collection statistics are shown in Table S2.

Four-wavelength anomalous dispersion analysis of the mercury derivative (space group $P\,4\,2_1\,2$) was performed using CRANK2 (24), followed by iterative density modification and automated model building using parrot (25) and buccaneer (26, 27), part of the CCP4 program suite (28). An initial model comprising a single pUL7:pUL51(8–142) core heterodimer was used as a molecular replacement model to solve the structure of the native complex (space group $P\,2_1$) using MolRep (29), identifying four core heterodimers in the asymmetric unit with *pseudo* four-fold non-crystallographic symmetry. Density modification and automated model building were performed using parrot and buccaneer, respectively, followed by cycles of iterative manual rebuilding in COOT (30) and TLS plus positional refinement using Refmac5 (31) with local NCS restraints. The building was assisted by the use of real-time molecular dynamics-assisted model building and map fitting with the program ISOLDE (32). Final cycles of refinement following manual rebuilding were performed using autoBUSTER (33) with local NCS restraints and TLS groups that were identified with the assistance of the TLSMD server (34). The quality of the model was monitored throughout the refinement process using Molprobity (35) and the validation tools in COOT. Molecular graphics were produced using PyMOL (36). Conservation of pUL7 residues across the α-herpesviruses was mapped onto the structure using the CONSURF server (37) and the sequence alignment used for co-evolutionary analysis (*Data set 2*, below).

*Bioinformatics and evolutionary analysis*

Protein sequences of pUL7 and pUL51 homologues from representative α-, β- and γ-herpesviruses that infect humans were as follows (Uniprot ID): HSV-1 pUL7 (A0A110B4Q7) and pUL51 (D3YPL0), VZV pORF53 (P09301) and pORF7 (P09271), HCMV pUL103 (D3YS25) and pUL71 (D3YRZ9), HHV-7 U75 (P52458) and U44 (P52474), KSHV pORF42 (F5HAI6) and pORF55 (F5H9W9), EBV BBRF2 (P29882) and BSRF1 (P0CK49). Secondary structure prediction was performed using the NetSurfP-1.1 server (38), disorder prediction was performed using moreRONN version 4.6 (39) and palmitoylation sites were predicted using CSS-Palm 4.0 (40) using the confidence threshold 'High'. Structure-based database searches for proteins with similar folds to pUL7 or pUL51 were performed using PDBeFOLD (41), DALI (42) and CATHEDRAL (43).

Clustal Omega (44) was used to generate seed alignments for *Alphaherpesvirinae* (HSV-1, VZV) or across all sub-families (HSV-1, VZV, HCMV, HHV7, KSHV, EBV). Seed alignments were used to generate hidden Markov models (HMMs) using the HMMER (45) program hmmbuild. HMMs were subsequently used to extract and align homologue sequences from UniProt using HMMER (45) program hmmsearch locally (for *Alphaherpesvirinae*) or using the HMMER web server (46) (for all *Herpesviridae*). We mapped the proteins thus identified to the source virus genomes, discarding any protein sequences from partial genome sequences where pUL7 or pUL51 were absent. Our initial alignments comprised distinct pairs of pUL7 and pUL51 sequences from 205 *Alphaherpesvirinae*, 147 *Betaherpesvirinae* and 78 *Gammaherpesvirinae*, and the alignments for homologues in each subfamily were improved by manual correction.

The structure of the core pUL7:pUL51 heterodimer was inspected to compile a table of 63 pairwise interactions between amino acids in the two proteins, 59 of which involved side chain atoms. These interactions arose from 33 distinct residues in pUL7 and 29 residues in pUL51. Using the alignments generated above, we compiled a matrix of amino acid pairs (one in each pUL7 and pUL51 homologue) that are predicted to interact. For each pair of interacting sites, we calculated the strength of the correlation between its amino acid states across the alignment. For this purpose, we followed Zaykin and colleagues (equation 3 of (47)). For a single pair of sites, whose alignments contain, respectively, *k* and *m* amino acid states, then the correlation between two of those states, *i* and *j*, is

$$r_{ij} = \frac{p_{ij} - p_i p_j}{\sqrt{p_i(1-p_i)}\sqrt{p_j(1-p_j)}}$$

where $p_i$ is the proportion of strains that carry amino acid *i* at the relevant site in pUL7, $p_j$ is the proportion that carry amino acid *j* in pUL51, and $p_{ij}$ is the proportion of strains that carry both. The total strength of correlation at the pair, *T*, is

$$T = \frac{(k-1)(m-1)}{km} N \sum_{i=1}^{k} \sum_{j=1}^{m} r_{ij}^2$$

where *N* is the number of strains, and the test statistic, *z* is this quantity summed across all interacting pairs.

$$z = \sum_{i=1}^{I} T$$

where *I* is the number of interactions. To test whether *z*, the signature of coevolution, was significantly greater than would be expected by chance, we compared the measured test statistic to a null distribution comprised of $10^6$ data sets for which the interacting partner sites were randomly permuted. The *p* value for each test was the proportion of randomly permuted data sets for which the test statistic was greater than or equal to the value for the real data. Under our permutation scheme, each randomized data set resembled the true data in terms of the total number of interactions, the number of interactions involving each site, and the allele frequencies at each putatively interacting site. The test also controls for shared evolutionary history, which can generate spurious evidence of coevolution (48). As a consequence, however, the test is expected to be highly conservative, because many of the randomized interactions might resemble the true interactions (not least because single sites were involved in multiple putative interactions) and because, under plausible evolutionary scenarios, multiple interacting pairs might evolve in concert.

Of this set of interactions, not all could be analyzed for all sequences, either because of missing amino acids in some sequences (due to both deletions and missing data), or because we could not be confident of the alignment of some sites. There was thus an inherent trade-off between maximizing the number of interactions and maximizing the number of strains in the test. We initially examined alignments across the *Herpesviridae*, but the low sequence identity meant that we could not confidently assign homology for most sites involved in putative interactions. Across the family as a whole, only 12 conserved interacting pairs could be analyzed, and this led to an underpowered test. Accordingly, we restricted our analyses to the *Alphaherpesvirinae*. From our initial alignments we excluded six very short sequences (one pUL51 homologue: A0A2Z4H851, and five pUL7 homologue: A0A120I2R6, A0A097HXP5, A0A286MM74, A0A2Z4H5E9, A0A120I2N0). This led to an alignment containing 199 strains, for which the amino acids of 35/63 interacting sites could be confidently aligned across all strains. These 35 interactions involved 21 sites from pUL51 and 19 sites from pUL7 (*Data set 1*, Table S4, and main text).

Because so many interactions were missing from this analysis, we next excluded two further pUL7 homologue sequences (B7FEJ7, A0A0X8E9M8) where many of the interacting sites could not be confidently aligned. This led to an alignment of 197 strains, for which 54/63 interactions could be tested (involving all 29 putatively interacting sites from pUL51 and 29/33 sites from pUL7). Despite the increase in the size of the data set, results were little changed (*Data set 2*, Table S4). Results were similarly little changed when we considered only interactions involving side chain atoms (*Data set 3*, Table S4), and when restricted our analysis to the subset of better conserved positions, as found in the regions of aligned sequence returned by HMMER (*Data set 4*, Table S4).

*Mammalian cell culture and transfection*

Mycoplasma-free human embryonic kidney (HEK) 293T and HeLa cells were maintained in Dulbecco's modified Eagle's medium (DMEM; ThermoFisher) supplemented with 10% (v/v) heat-inactivated fetal bovine serum (FBS) and 2 mM L-glutamine (ThermoFisher). Cells were maintained at 37°C in a humidified 5% $CO_2$ atmosphere.

Plasmids for GFP-pUL7 (N-terminal tag) and pUL51-mCherry (C-terminal tag) were as used in (1). Homologues from other herpesviruses were cloned into pEGFP-C2, encoding an N-terminal GFP tag, or pmCherry-N1, encoding a C-terminal mCherry tag, as follows. pUL103 and pUL71 were cloned from HCMV strain Toledo cDNA, pORF42 and pORF55 were cloned from KSHV strain JSC-1 cDNA, and VZV pORF53 and pORF7 were cloned from codon-optimized synthetic genes (GeneArt) to boost their otherwise-poor expression in cultured cells.

For co-precipitation experiments, $5\times10^6$ HEK 293T cells were transfected by adding 1 µg total DNA (split evenly by mass between the plasmids indicated) and 1.5 µg of branched polyethylenimine (PEI; average MW ~25,000, Merck) that had been diluted in Opti-MEM (ThermoFisher) and incubated together for 20 min before addition to cells.

For immunocytochemistry, $7.5\times10^4$ HeLa cells/well were seeded in six-well plates containing four sterile no. 1.5 coverslips/well and grown overnight before being transfected by addition of 625 ng total DNA (split evenly by mass between the plasmids indicated) and 6 µL/well TransIT-LT1 (Mirus) that had been diluted in Opti-MEM and incubated together for 20 min before addition to cells.

*Co-precipitation and immunoblotting*

Cells were harvested 24 h post-transfection by scraping in phosphate buffered saline (PBS; 137 mM NaCl, 2.7 mM KCl, 10 mM $Na_2HPO_4$, 1.8 mM $KH_2PO_4$), and washed twice in PBS. Cell pellets were resuspended in lysis buffer (10 mM tris pH 7.5, 150 mM NaCl, 0.5 mM EDTA, 0.5% IGEPAL CA-630 (a.k.a. NP-40, Merck), 1% (v/v) EDTA-free protease inhibitor cocktail (Merck)) and incubated at 4°C for 30 min before clarification by centrifugation at 20,000×g, 4°C for 10 min. The protein concentration in each lysate was normalized after assessment using the bicinchoninic acid assay (ThermoFisher) according to the manufacturer's instructions. Normalised lysates were incubated for 1 h at 4°C with GFP-Trap or RFP-Trap bead slurry (Chromotek) that had been pre-equilibrated in wash buffer (10 mM tris pH 7.5, 150 mM NaCl, 0.5 mM EDTA). Following incubation, beads were washed three times, the supernatant was completely removed, beads were resuspended in SDS-PAGE loading buffer and the samples were heated at 95 °C for 5 min to liberate bound proteins before removal of the beads by centrifugation. Samples were separated by SDS-PAGE using 12% or 15% polyacrylamide gels and transferred to Protran nitrocellulose membranes (Perkin Elmer) using the Mini-PROTEAN and Mini-Trans-

Blot systems (BioRad) following the manufacturer's protocol. After blocking in PBS with 5% (w/v) non-fat milk powder, membranes were incubated with primary antibody overnight at 4°C and then secondary antibody for 1 h at room temperature. Dried blots were visualized on an Odyssey CLx infrared scanner (LI-COR).

*Immunocytochemistry*

Cells were transferred onto ice 24 h post-transfection. Coverslips were washed with ice-cold PBS and incubated with cold 20 mM HEPES pH 7.5, 4% (v/v) electron microscopy-grade formaldehyde (PFA, Polysciences) for 5 minutes on ice before being incubated with 20 mM HEPES pH 7.5, 8% (v/v) PFA at room temperature for 10 minutes. Coverslips were washed with PBS before quenching of residual PFA by addition of 25 mM $NH_4Cl$ for 5 min at room temperature. After washing with PBS, cells were permeabilized by incubation with 0.1% saponin in PBS for 30 min before being incubated with blocking buffer (5% (v/v) FBS, 0.1% saponin in PBS) for 30 min. Primary antibodies (below) were diluted in blocking buffer and incubated with coverslips for 2 h. Coverslips were washed five times with blocking buffer before incubation for 1 h with the relevant secondary antibodies (below) diluted in blocking buffer. Coverslips were washed five times with blocking buffer, three times with 0.1% saponin in PBS, three times with PBS, and finally with ultrapure water. Coverslips were mounted using Mowiol 4-88 (Merck) containing 200 nM 4′,6-diamidino-2-phenylindole (DAPI) and allowed to set overnight. Images were acquired using a Zeiss LSM780 confocal laser scanning microscopy system mounted on an AxioObserver.Z1 inverted microscope using a 64× Plan Apochromat objective (NA 1.4). Images were processed using Fiji (49, 50).

*Antibodies*

Primary antibodies used for immunoblotting were rabbit anti-GFP (Merck, G1544), rat anti-RFP (Chromotek, 5F8), or mouse anti-GAPDH (ThermoFisher, AM4300). Secondary antibodies for immunoblotting were LI-COR IRDye 680T conjugated goat anti-rat (926-68029), donkey anti-rabbit (926-68023) or goat anti-mouse (926-68020), or LI-COR IRDye 800CW conjugated donkey anti-rabbit (926-32213) or goat anti-mouse (926-32210). Primary antibodies used for immunocytochemistry were anti-TGN46 (Bio-Rad, AHP500G), mouse anti-Paxillin (BD Biosciences 610051), rabbit anti-Zyxin (abcam ab71842), and secondary antibodies were Alexa Fluor 647 conjugated donkey anti-sheep (A-21448, ThermoFisher), goat anti-mouse (A-21236, ThermoFisher) or goat anti-rabbit (A-21245, ThermoFisher).

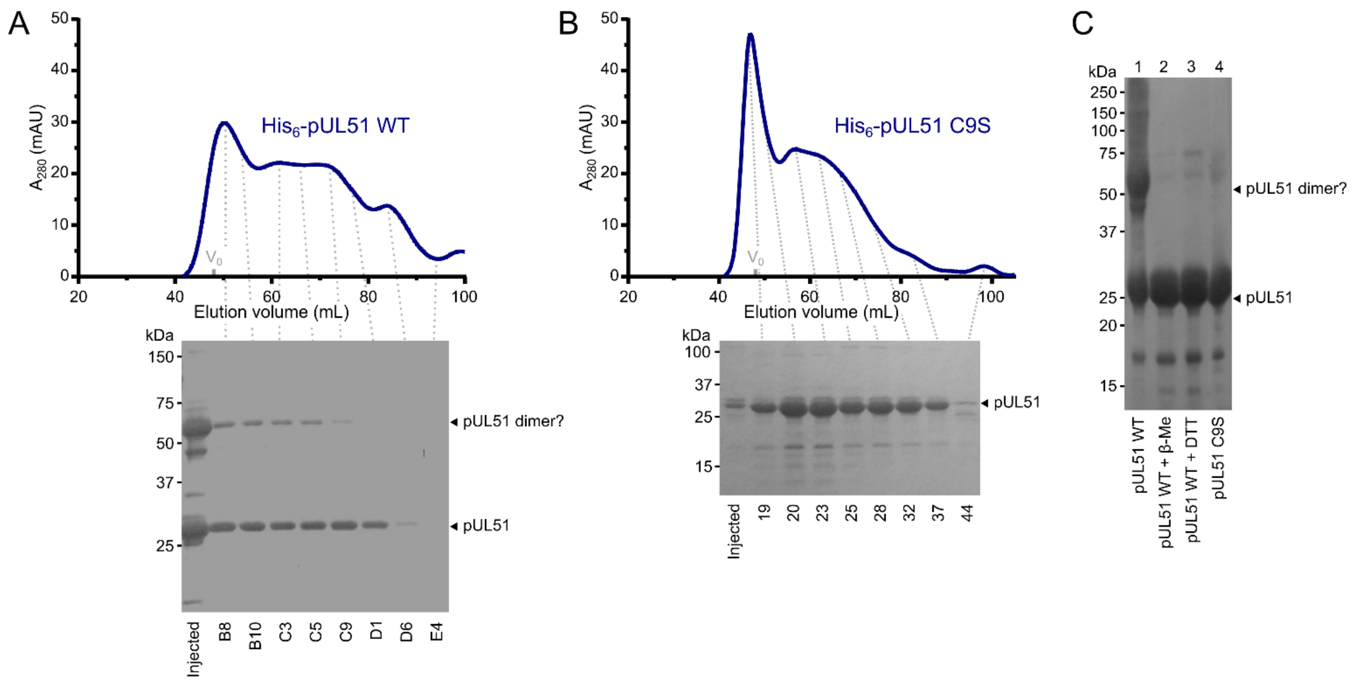


**Figure S1. HSV-1 pUL51 forms large soluble aggregates when purified in isolation.** (A) SEC elution profiles of His6-tagged wild-type pUL51 (A) and His6-tagged pUL51 C9S (B). Proteins were injected onto an S200 16/600 column (GE Healthcare) equilibrated in 20 mM Tris (pH 7.5), 200 mM NaCl, 1 mM DTT. Both proteins have extended elution profiles with peaks near the column void volume ($V_0$), consistent with their forming large soluble aggregates. Coomassie-stained SDS-PAGE analysis of eluted SEC fractions are shown beneath each chromatogram. Note that there is a higher molecular weight band in (A), consistent with the presence of an SDS-resistant pUL51 dimer, despite the presence of 1 mM DTT in the SEC buffer and 2 mM DTT in the SDS-PAGE loading buffer. (C) Purified His6-tagged wild-type pUL51 was subjected to SDS-PAGE either without additional treatment (*lane 1*) or following incubation with 50 mM β-mercaptoethanol (*lane 2*) or 50 mM DTT (*lane 3*). Comparison with the His6-tagged pUL51 C9S mutant (*lane 4*) confirms that cysteine 9, the residue that becomes palmitoylated in mammalian cells (51), mediates disulfide bond mediated dimerization of recombinant wild-type pUL51. C9S substituted pUL51 (or truncations thereof) was thus used for all subsequent experiments.

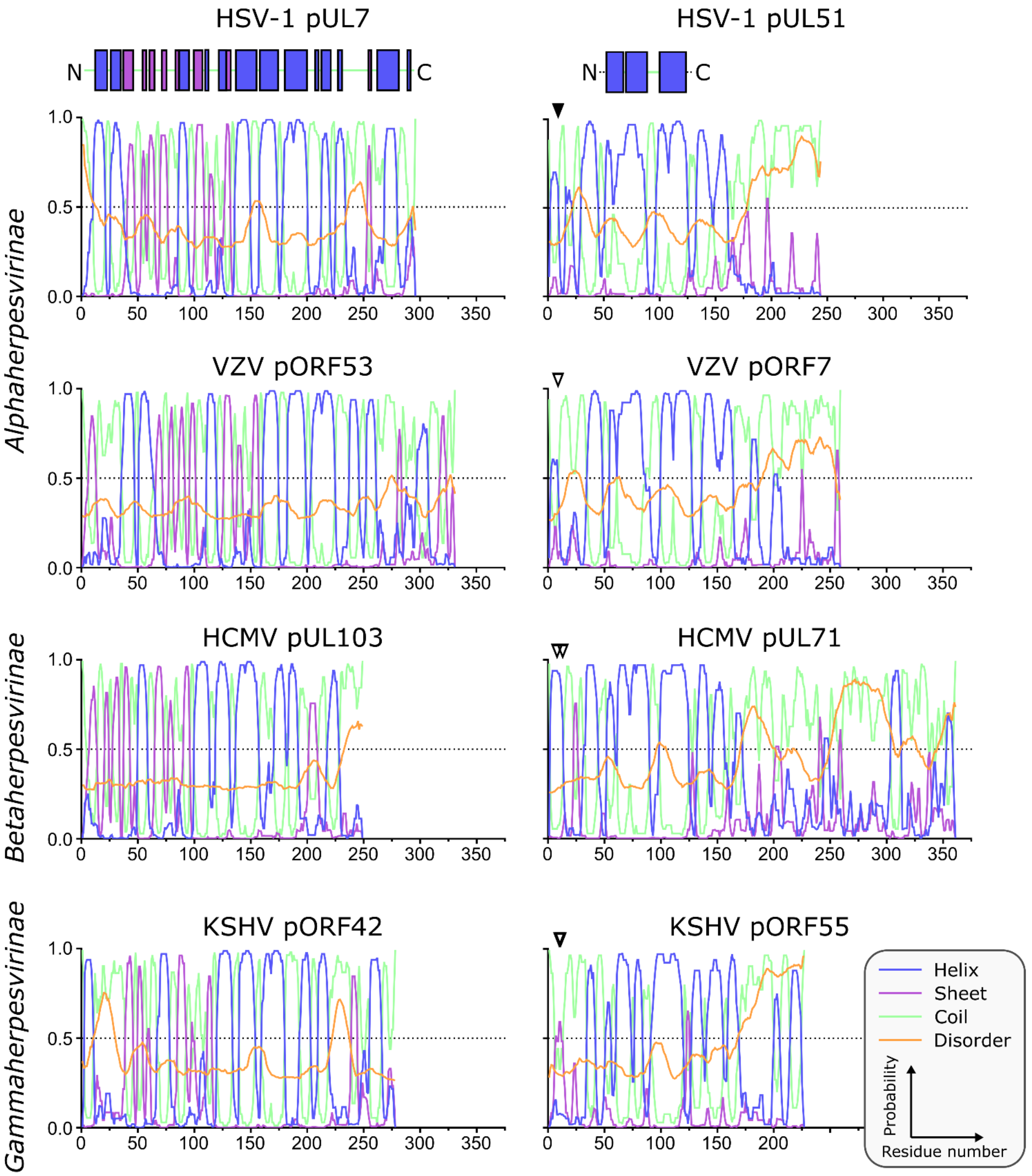


**Figure S2. Predicted secondary structure of pUL7 and pUL51 homologues from representative human α-, β- and γ-herpesviruses.** Analyses of amino acid sequences were performed as described in *SI Materials and Methods*. Per-residue probabilities of forming α-helix (blue), β-sheet (purple) or coil (green) are shown, as is the probability of disorder (orange). Residues that are known (solid triangles) or predicted (empty triangles) to be palmitoylated are marked: Cys9 of HSV-1 pUL51, Cys9 of VZV pORF7, Cys8 and Cys13 of HCMV pUL71, Cys10 and Cys11 of KSHV pORF55. Regions of pUL7 and pUL51 α-helix and β-sheet observed in the pUL7:pUL51(8–142) core heterodimer structure are shown above the predictions as boxes. The predicted pUL7 and pUL51 secondary structural elements are largely conserved across herpesvirus families, although the first two helices of pUL7 are not conserved in β-herpesviruses like HCMV. Additionally, the C-terminal regions of pUL51 homologues vary in length, although in all herpesvirus families they are predicted to be largely unstructured.

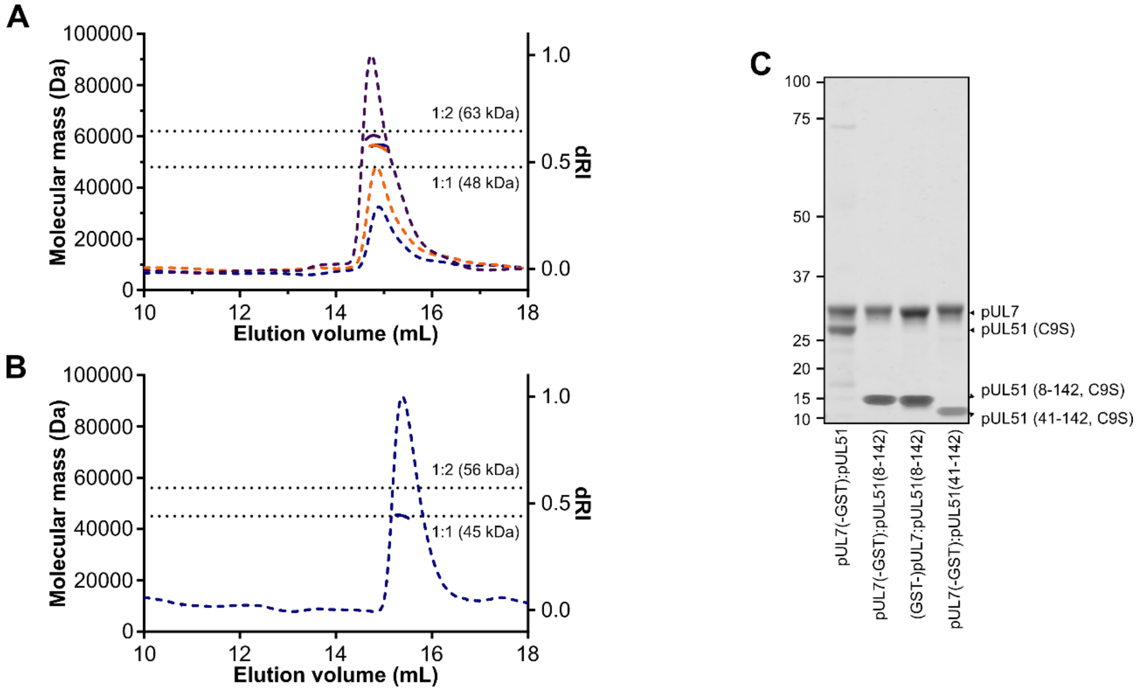


**Figure S3. SEC-MALS of truncated pUL7:pUL51 complexes.** SEC elution profiles (differential refractive index, dashed lines) and weight-averaged molecular masses across the elution peaks (solid lines) are shown. (A) SEC-MALS of pUL7:pUL51(8-142) where pUL7 had been purified using an N-terminal GST tag that was subsequently removed using human rhinovirus 3C protease. Observed mass for the pUL7:pUL51(8-142) complex was 57.4 ± 2.2 kDa, compared with a theoretical mass of 62.7 kDa for a 1:2 heterotrimer. Samples were injected onto the column at 0.3 mg/mL (blue), 0.5 mg/mL (orange) and 1 mg/mL (purple). (B) SEC-MALS of pUL7:pUL51(41-142), injected onto the column at 0.3 mg/mL. The observed mass was 45.1 kDa, compared to a theoretical mass of 44.8 kDa for a 1:1 heterodimer (C) Coomassie-stained SDS-PAGE analysis of samples used for SEC-MALS analysis in (A), (B) and Fig. 2. The GST purification tag was cleaved from all samples used for SEC-MALS, the pUL7 protein having been tagged at the N or C terminus during the initial purification steps as shown.

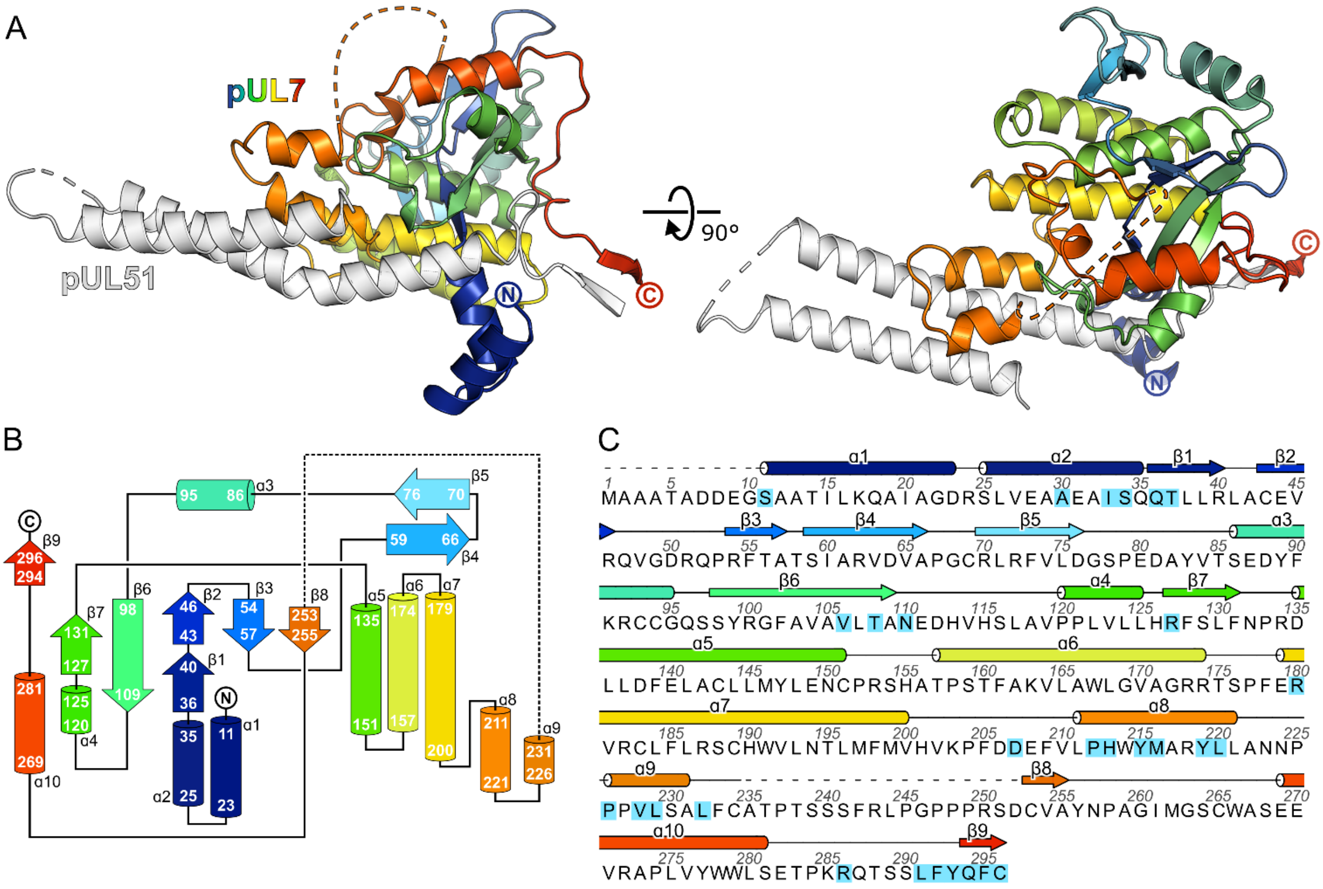


**Figure S4. The CUSTARD fold of pUL7.** (A) Structure of pUL7:pUL51(8–142) core heterodimer is shown as ribbons, with pUL51 colored white and pUL7 colored from blue (residue 11) to red (residue 296). Two orthogonal views are shown. (B) Schematic diagram of the topology of pUL7. (C) HSV-1 pUL7 sequence, with secondary structure shown above. Residues that interact with pUL51 are highlighted in cyan.

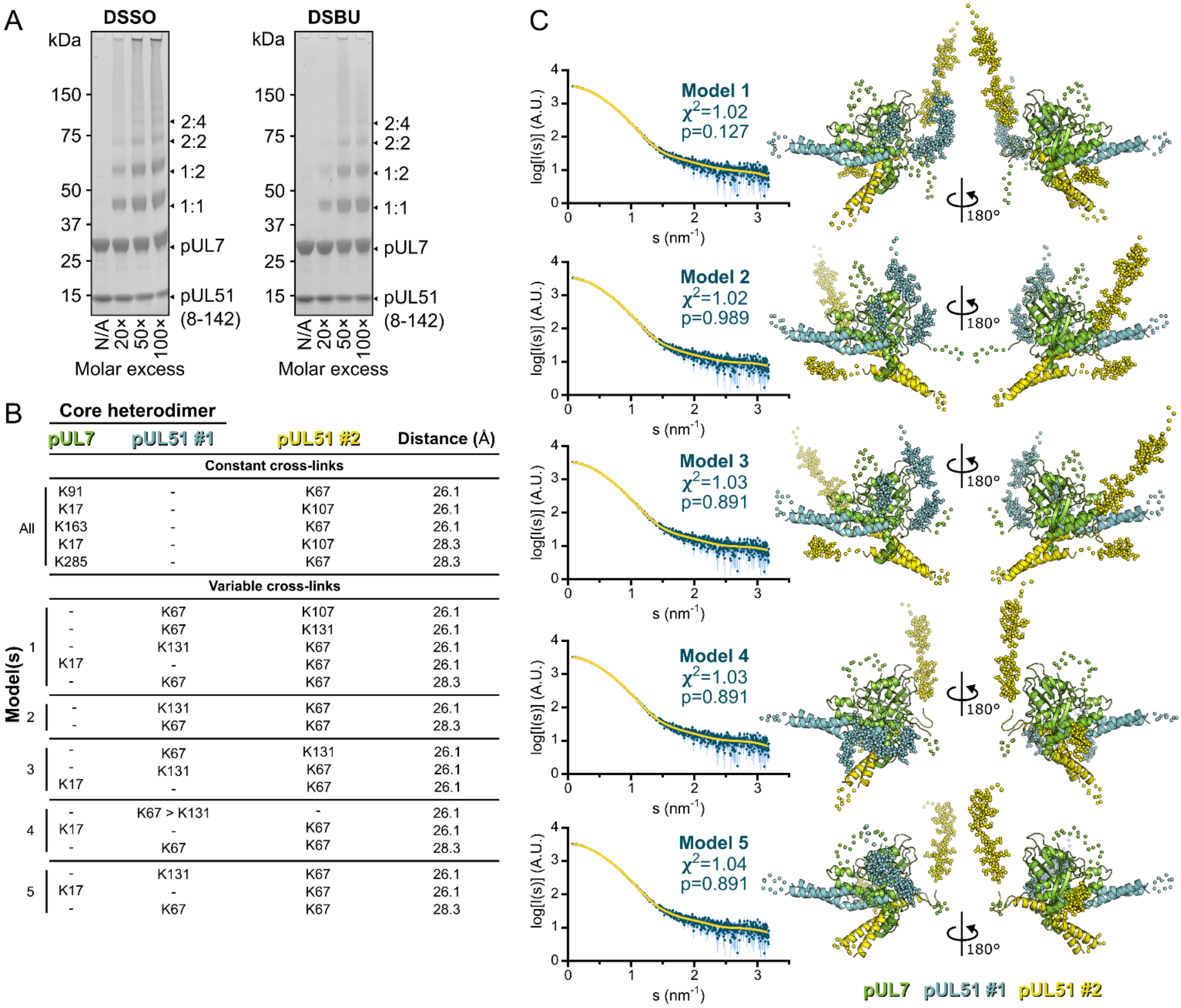


| Core heterodimer | pUL7 | pUL51 #1 | pUL51 #2 | Distance (Å) |
|---|---|---|---|---|
| **Constant cross-links** | | | | |
| All | K91 | - | K67 | 26.1 |
| | K17 | - | K107 | 26.1 |
| | K163 | - | K67 | 26.1 |
| | K17 | - | K107 | 28.3 |
| | K285 | - | K67 | 28.3 |
| **Variable cross-links** | | | | |
| Model(s) 1 | - | K67 | K107 | 26.1 |
| | - | K67 | K131 | 26.1 |
| | - | K131 | K67 | 26.1 |
| | K17 | - | K67 | 26.1 |
| | - | K67 | K67 | 28.3 |
| 2 | - | K131 | K67 | 26.1 |
| | - | K67 | K67 | 28.3 |
| 3 | - | K67 | K131 | 26.1 |
| | - | K131 | K67 | 26.1 |
| | K17 | - | K67 | 26.1 |
| 4 | - | K67 > K131 | - | 26.1 |
| | K17 | - | K67 | 26.1 |
| | - | K67 | K67 | 28.3 |
| 5 | - | K131 | K67 | 26.1 |
| | K17 | - | K67 | 26.1 |
| | - | K67 | K67 | 28.3 |

**Figure S5. Cross-linking mass spectrometry analysis and pseudo-atomic modelling of the pUL7:pUL51(8–142) solution heterotrimer.** (A) Coomassie-stained SDS-PAGE analysis of purified pUL7:pUL51(8–142) following 30 min incubation at room temperature with varying molar excesses of the cross-linking agents DSSO (left) or DSBU (right). Theoretical migration of proteins corresponding to pUL7, pUL51(8–142), and 1:1, 1:2, 2:2 or 2:4 complexes thereof, are indicated. (B) Cross-linking restraints used for pseudo-atomic modelling of the pUL7:pUL51(8–142) heterotrimer. Restraints used for all models (“constant cross-links”) and permuted restraints (“variable cross-links”) that were used for the five best-fit (lowest $\chi^2$) models are shown. (C) The five best pseudo-atomic models (lowest $\chi^2$) generated by fitting to the pUL7:pUL51(8–142) SAXS profile as described in *SI Materials and Methods* using the restrains shown in (B). The core heterodimer of pUL7 (residues 11–234 and 253–296; green) and pUL51 (residues 41–89 and 96–125; #1, cyan) and the additional molecule of pUL51 (residues 41–89 and 96–125; #2, yellow) are shown as ribbons (right). Additional regions modelled using I-TASSER or CORAL are shown as spheres. The fit of the computed scattering (yellow) to the pUL7:pUL51(8–142) SAXS profile (aqua) is shown for each model (left), as are reduced $\chi^2$ and CorMap *P*-values (7, 9).

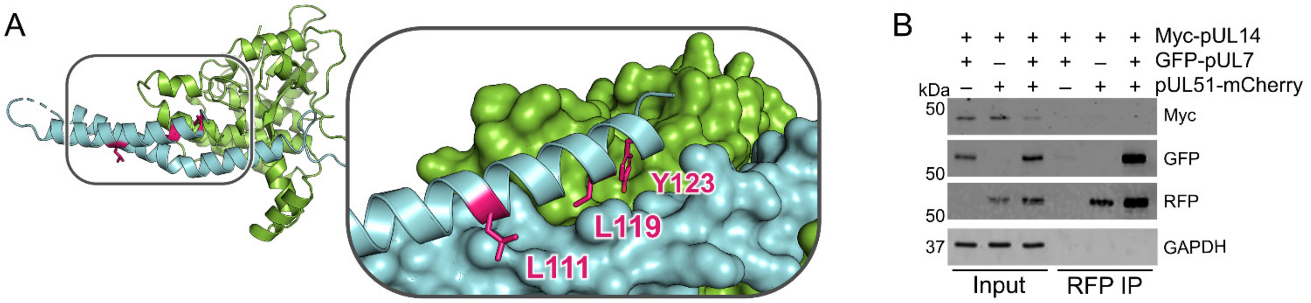


**Figure S6. pUL51 does not co-precipitate pUL14 in uninfected cultured cells.** (A) Core heterodimer of pUL7:pUL51 with residues required for the reported pUL51 and pUL14 (52) highlighted in pink. Inset shows pUL7 and the first helix of pUL51 as surfaces, and side chains of residues that were substituted with alanine in (52) are shown as sticks. (B) HEK 293T cells were co-transfected with myc-tagged pUL14 from HSV-1 along with GFP-pUL7 and pUL51-mCherry, as shown. Co-immunoprecipitation of myc-pUL14 with pUL51-mCherry is not observed either in the presence or absence of pUL7.

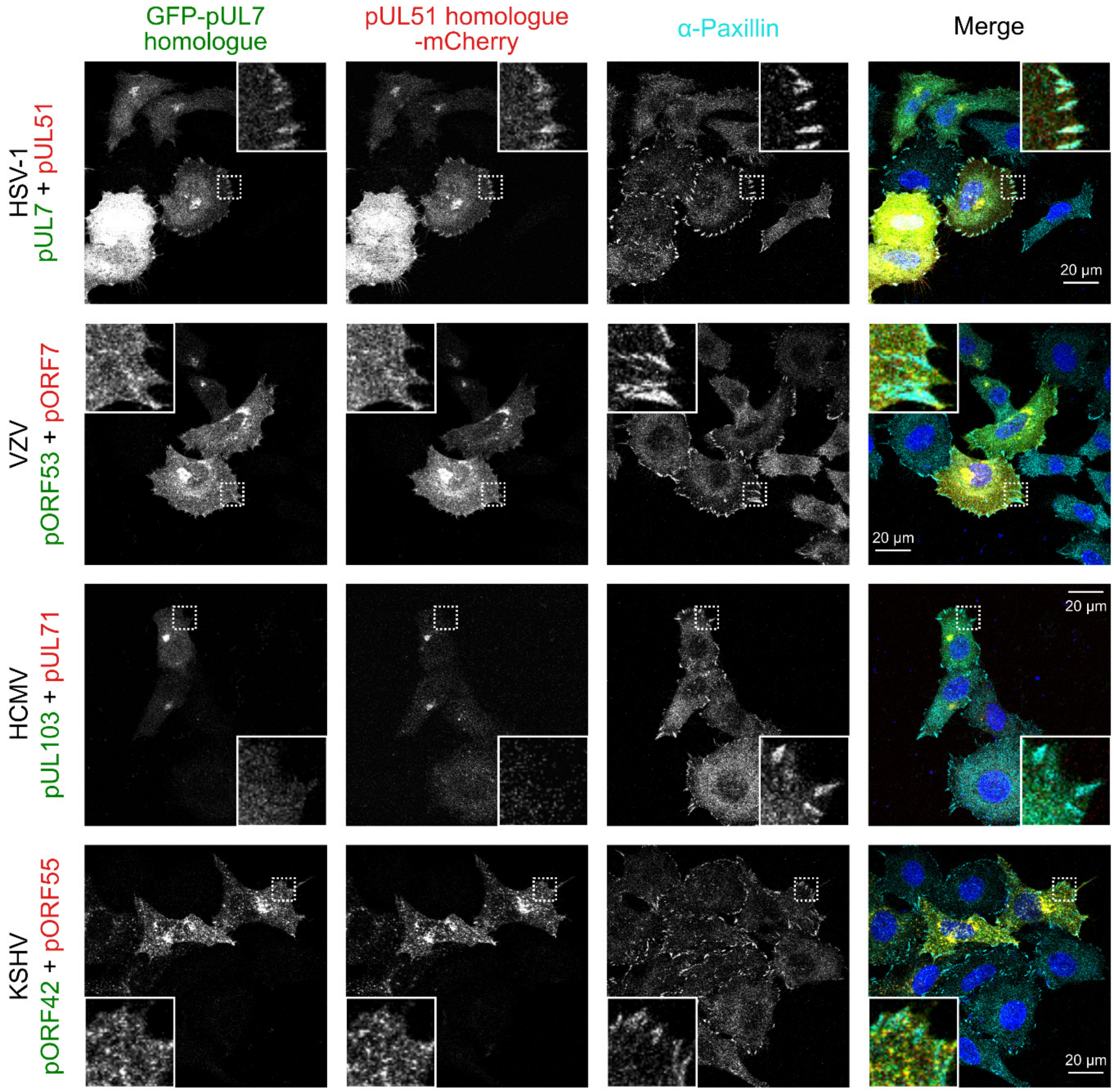


**Figure S7. The pUL7:pUL51 complex from HSV-1 co-localizes with focal adhesion marker paxillin, but homologues from other human herpesviruses do not.** HeLa cells were co-transfected with GFP-pUL7 and pUL51-mCherry, or with similarly-tagged homologues from VZV, HCMV and KSHV. Cells were fixed 24 hours post-transfection and immunostained for the focal adhesion marker protein paxillin before imaging by confocal microscopy. Co-localization between the GFP, mCherry and far-red (paxillin) fluorescence was only observed for cells transfected with HSV-1 pUL7-GFP and pUL51-mCherry.

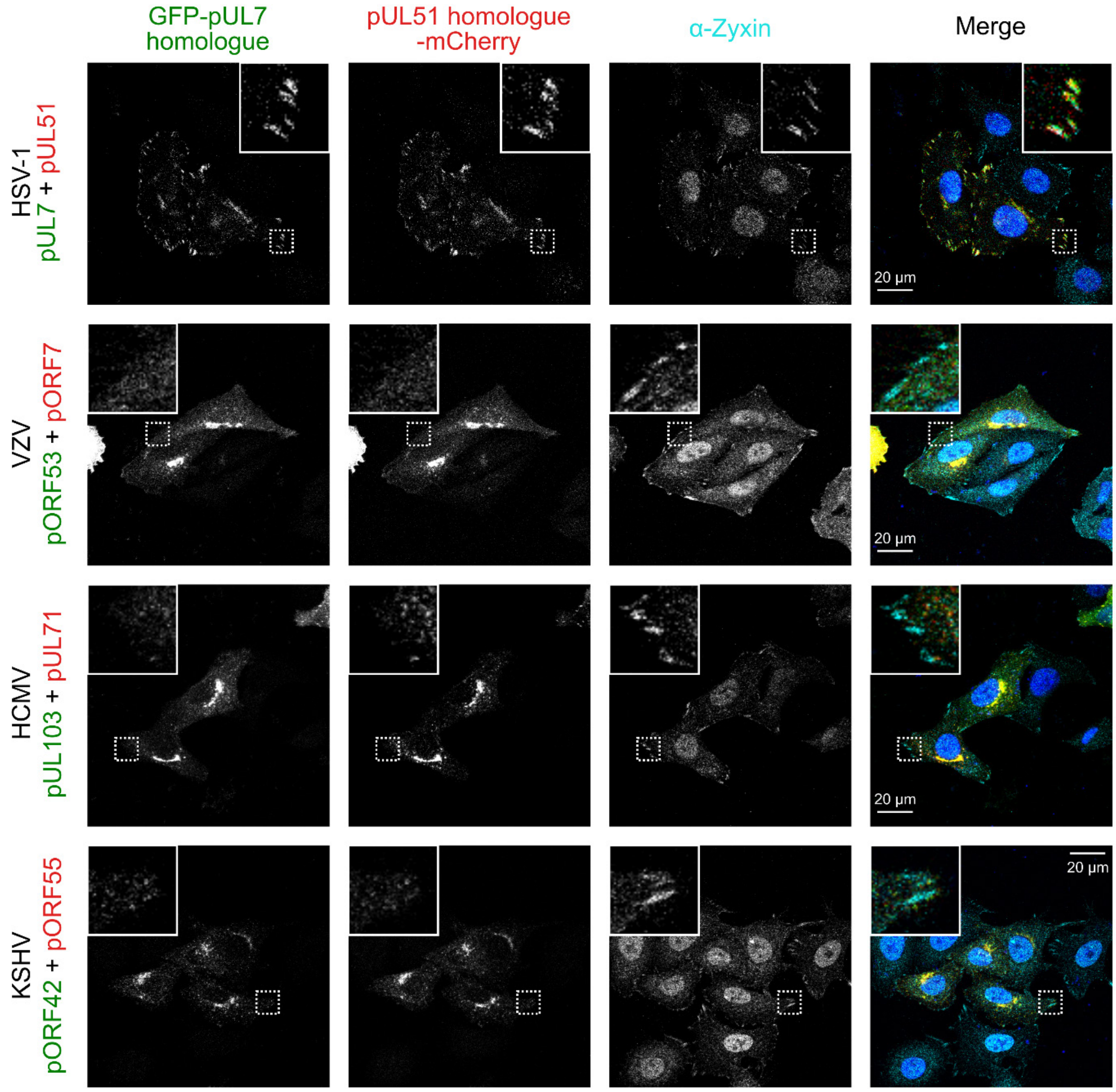


**Figure S8. The pUL7:pUL51 complex from HSV-1 co-localizes with focal adhesion marker zyxin but homologues from other human herpesviruses do not.** HeLa cells were co-transfected with GFP-pUL7 and pUL51-mCherry, or with similarly-tagged homologues from VZV, HCMV and KSHV. Cells were fixed 24 hours post-transfection and immunostained for the focal adhesion marker protein zyxin before imaging by confocal microscopy. Co-localization between the GFP, mCherry and far-red (zyxin) fluorescence was only observed for cells transfected with HSV-1 pUL7-GFP and pUL51-mCherry.

**Table S1. SAXS data collection and analysis parameters**. Program version numbers are shown in parentheses.

| | pUL7:pUL51 | | pUL7:pUL51 (8-142) |
|---|---|---|---|
| | 1:2 complex | 2:4 complex | 1:2 complex |
| *Data-collection parameters* | | | |
| Radiation Source | Petra III (DESY, Hamburg, Germany) | | |
| Beamline | EMBL P12 | | |
| Detector | Dectris Pilatus 6M | | |
| Beam geometry (mm) | 0.2 × 0.12 | | |
| X-ray wavelength (nm) | 0.124 | | |
| Sample-to-detector distance (m) | 3.0 | | |
| Temperature (°C) | 20.2 | | |
| Measured *s*-range ($nm^{-1}$) | 0.0225–7.299 | | |
| Exposure time (s) | 0.995 | | |
| Injected protein concentration (mg $ml^{-1}$) | 8 | | 4.5 |
| Shannon-channel limited *s*-range ($nm^{-1}$) | < 5.57 | < 6.70 | < 7.16 |
| *Structural parameters* | | | |
| $I(0)$ (a.u.*) [from $p(r)$] | 4624 ± 13 | 3136 ± 11 | 3098 ± 5 |
| Real-space $R$ g (nm) [from $p(r)$] | 4.30 ± 0.03 | 4.79 ± 0.04 | 3.0 ± 0.01 |
| $I(0)$ (a.u.*) (from Guinier) | 4538 ± 10 | 3100 ± 7 | 3331 ± 2 |
| $R_g$ (nm) (from Guinier) | 3.95 ± 0.07 | 4.56 ± 0.28 | 2.96 ± 0.01 |
| $D_{max}$ (nm) | 18.2 | 19.7 | 11.5 |
| Porod volume estimate (Vp, $nm^3$) | 160 | 340 | 116 |
| *Molecular-mass determination* | | | |
| Molecular mass $M_r$, kDa [from SAXSMOW] | 95 | 177 | 73 |
| Molecular mass $M_r$, kDa [from Vc] | 81 | 167 | 66 |
| Molecular mass $M_r$, kDa [from Bayesian consensus] | 86–96 | 162–195 | 66–73 |
| Expected $M_r$ from sequence, kDa | 84.37 | 168.7 | 63.11 |
| *Software employed* | | | |
| Primary data reduction | *CHROMIXS* | | |
| Data processing | *PrimusQT/GNOM(5.0)* | | |
| *Ab initio* analysis | *DAMMIN(5.3)/GASBOR(2.3i)* | | |
| Spatial averaging and resolution estimates | *DAMAVER(5.0)* | | |
| Computation of model intensities | *CRYSOL(2.8.3)* | | |
| Pseudo-atomic modelling | *CORAL(1.1)* | | |
| *Small Angle Scattering Biological Data Bank* | | | |
| SASBDB accession codes | SASDG57 | SASDG47 | SASDG37 |

*arbitrary units

**Table S2. Crystallographic data collection and refinement statistics**. Statistics for the highest-resolution shell are shown in parentheses.

| | | Mercury(II) acetate derivative | | | |
|---|---|---|---|---|---|
| | Native | Remote (High E) | Peak | Inflection | Remote (Low E) |
| *Data collection* | | | | | |
| Wavelength (Å) | 0.97625 | 0.99702 | 1.00728 | 1.00809 | 1.01627 |
| Space group | $P\,2_1$ | $P\,4\,2_1\,2$ | | | |
| Cell dimensions | | | | | |
| *a, b, c* (Å) | 79.51, 106.3, 106.0 | 106.5, 106.5, 79.0 | 106.5, 106.5, 79.2 | 106.5, 106.5, 79.3 | 106.5, 106.5, 79.4 |
| *α, β, γ* (°) | 90, 92.0, 90 | 90, 90, 90 | 90, 90, 90 | 90, 90, 90 | 90, 90, 90 |
| Resolution (Å) | 105.9–1.8 (1.86–1.83) | 53.2–2.8 (2.96–2.80) | 53.2–2.9 (3.02–2.87) | 53.3–2.9 (3.09–2.91) | 53.3–3.0 (3.16–2.98) |
| Unique reflections | 154,984 (7654) | 11,618 (1627) | 10,931 (1561) | 10,498 (1643) | 9789 (1515) |
| Completeness (%) | 100 (99.9) | 99.7 (97.9) | 99.9 (99.5) | 99.9 (99.7) | 99.7 (98.5) |
| Anomalous | – | 99.7 (97.9) | 99.9 (99.5) | 99.9 (99.7) | 99.7 (98.5) |
| Multiplicity | 6.5 (6.5) | 47.6 (36) | 24.9 (25.6) | 24.8 (24.9) | 24.7 (24.8) |
| Anomalous | – | 25.9 (19.1) | 13.5 (13.5) | 13.5 (13.2) | 13.5 (13.2) |
| $R_{merge}$ | 0.120 (3.197) | 0.261 (2.664) | 0.231 (2.506) | 0.235 (2.348) | 0.232 (2.158) |
| $R_{pim}$ | 0.051 (1.367) | 0.038 (0.435) | 0.047 (0.502) | 0.048 (0.476) | 0.047 (0.439) |
| $CC_{1/2}$ | 0.996 (0.321) | 0.999 (0.804) | 0.999 (0.783) | 0.999 (0.792) | 0.999 (0.809) |
| $CC_{anom}$ | – | 0.830 (0.059) | 0.741 (0.000) | 0.664 (0.014) | 0.224 (0.000) |
| Mean I/σ(I) | 6.8 (0.5) | 14.3 (1.7) | 12.1 (1.6) | 12.4 (1.7) | 12.4 (1.8) |
| Wilson B ($A^2$) | 35.2 | | | | |
| *Refinement* | | | | | |
| Resolution (Å) | 26.6–1.8 (1.84–1.83) | | | | |
| Reflections | | | | | |
| Working set | 154,885 (2892) | | | | |
| Test set | 8093 (206) | | | | |
| $R_{work}$ | 0.194 (0.240) | | | | |
| $R_{free}$ | 0.220 (0.244) | | | | |
| No. of atoms | | | | | |
| Protein | 11692 | | | | |
| Solvent | 717 | | | | |
| Other* | 28 | | | | |
| Root mean square deviation | | | | | |
| Bond lengths (Å) | 0.009 | | | | |
| Bond angles (°) | 0.90 | | | | |
| Ramachandran favoured (%) | 98.7 | | | | |
| Ramachandran outliers (%) | 0.0 | | | | |
| Poor rotamers (%) | 0.39 | | | | |
| Mean B value ($A^2$) | 45.2 | | | | |

*Glycerol molecules and chloride ions

**Table S3. pUL7:pUL51(8–142) cross-links identified by mass spectrometry.** Two protein bands from SDS-PAGE analysis, with molecular masses corresponding to 1:1 and 1:2 pUL7:pUL51(8–142), were analyzed.

| Reagent | Band | Protein | Residue | Sequence | Protein | Residue | Sequence |
|---|---|---|---|---|---|---|---|
| DSBU | 1:1 | pUL7 | 17 | AAATADDEGSA-ATIL[**K**]QAIAGDR | pUL51 | 107 | HHPGLEAPTIDG-AVAAHQD[**K**]MR |
| DSBU | 1:1 | pUL51 | 67 | RLV[**K**]AR | pUL51 | 67 | LV[**K**]AR |
| DSBU | 1:1 | pUL7 | 285 | APLVYWWLSETP[**K**]R | pUL51 | 67 | LV[**K**]AR |
| DSBU | 1:2 | pUL51 | 67 | RLV[**K**]AR | pUL51 | 67 | LV[**K**]AR |
| DSSO | 1:1 | pUL7 | 91 | FVLDGSPEDAYVTSEDYF[**K**]R | pUL51 | 67 | LV[**K**]AR |
| DSSO | 1:1 | pUL7 | 17 | AAATADDEGSA-ATIL[**K**]QAIAGDR | pUL51 | 107 | HHPGLEAPTIDG-AVAAHQD[**K**]MRR |
| DSSO | 1:1 | pUL51 | 107 | HHPGLEAPTIDGAVAAHQD[**K**]MR | pUL51 | 67 | RLV[**K**]AR |
| DSSO | 1:1 | pUL7 | 163 | SHATPSTFA[**K**]VLAWLGVAGR | pUL51 | 67 | LV[**K**]AR |
| DSSO | 1:1 | pUL51 | 131 | LADTCMATILQMYMSV-GAAD[**K**]SADVLVSQAIR | pUL51 | 67 | RLV[**K**]AR |
| DSSO | 1:1 | pUL7 | 17 | AAATADDEGSA-ATIL[**K**]QAIAGDR | pUL51 | 67 | LV[**K**]AR |

**Table S4. Co-evolution of the pUL7-pUL51 interaction interface across *Alphaherpesvirinae*.** $z$ is the sum of correlation values for interacting residue pairs and $p$ is the probability that this value would be expected by chance. The selection of sequences for each data set is described in *SI Materials and Methods*.

| | | | No. of interacting residues | | | |
|---|---|---|---|---|---|---|
| Dataset | No. strains, $N$ | No. interactions, $I$ | pUL51 | pUL7 | $z$ | $p$ |
| 1 | 199 | 35 | 21 | 19 | 4301.24 | 0.061 |
| 2 | 197 | 54 | 27 | 29 | 7319.44 | 0.066 |
| 3 | 197 | 52 | 26 | 29 | 6969.40 | 0.065 |
| 4 | 197 | 38 | 22 | 21 | 4559.47 | 0.062 |
